# A comprehensive view of somatic mosaicism by single-cell DNA analysis

**DOI:** 10.1101/2025.10.31.685648

**Authors:** Lovelace J. Luquette, Tim H. H. Coorens, Abhiram Natu, Milovan Suvakov, Ann Caplin, Mee Sook Jun, Alisa Mo, Joe Pelt, Lisa Anderson, Michele Berselli, Sravya Bhamidipati, Thomas Blanchard, Joseph Brew, Hye-Jung E. Chun, Hyunbin Chun, Mrunal K. Dehankar, William C. Feng, Rob Furatero, Christopher M. Grochowski, Eric Ho, Yeongjun Jang, Kavya Kottapalli, M. Kathryn Leonard, Nam Seop Lim, Tina Lindsay, Sarah Nicholson, Ivan Raimondi, Alexi Runnels, Constantijn Scharlee, JaeEun Shin, Alexander D. Veit, Melissa VonDran, Yifan Wang, Dennis J. Yuan, Yifan Zhao, SMaHT Single Cell Focus Group, Thomas J. Bell, Kristin Ardlie, Harsha Doddapaneni, Robert Fulton, Soren Germer, Dan Landau, Ji Won Oh, Peter J. Park, Flora M. Vaccarino, Christopher Walsh, Alexej Abyzov

## Abstract

Single-cell DNA sequencing offers a powerful means of studying somatic mosaicism but requires careful analysis to mitigate DNA amplification-related artifacts. We performed primary template-directed amplification (PTA) and sequencing of 102 nuclei from postmortem lung and colon tissues of a 74-year-old male. Single-cell mutation burdens and spectra were validated by duplex sequencing and revealed heterogeneity across organs and cells, including signatures of APOBEC activity and tobacco exposure. Cells from both tissues exhibited chromosomal aneuploidies, loss of chromosome Y, and chromosomal rearrangements including rearrangements of the T-cell receptor loci indicative of T-cells. Shared embryonic mutations between cells enabled reconstruction of cellular ancestries from the zygote, which were validated by bulk sequencing. Collectively, we demonstrate a comprehensive approach for single-cell genomics that yields an expansive view of diverse somatic mutation types from development through aging across diverse tissues—insights that are obscured in bulk sequencing and only partially captured by other single-cell methods.

## Introduction

Multiple studies spanning the past decade^1–7^ have converged into a coherent picture that every nucleated cell in the human body harbors a different genome—a phenomenon called somatic mosaicism. Somatic mosaicism can be studied in bulk tissue specimens that contain many cells, but this limits detection to the most frequent mutations within a tissue and obscures cellular heterogeneity. Although duplex sequencing methods^8–12^ have improved the accuracy of detection in bulk for small somatic mutations such as single nucleotide variants (SNVs) and small insertions and deletions (indels) by orders of magnitude, they primarily reveal average tissue mutational characteristics and have limited ability to identify large mutations such as structural variants (SVs).

The analysis of single cells—by *ex vivo* clonal expansion of individual cells^13–17^, laser capture microdissection (LCM) of small, naturally occurring clones^18–20^, or *in vitro* single-cell whole-genome amplification (scWGA)^5,6,21,22^ allows discovery of mutations in a cell independent of their frequency in bulk tissues. However, *ex vivo* expansion and LCM are limited to proliferating cells and scWGA has been challenging due to variable amplification uniformity across the genome^23,24^ and amplification-associated artifacts that resemble mutations. An important advance in scWGA was the development of primary template-directed amplification (PTA^25^), which greatly improved amplification uniformity and enabled detection of somatic SNVs with error rates of 1.3×10^-8^ per bp using carefully tailored computational methods^26^.

To date, PTA has been primarily applied to the postmortem human brain^21,22,26–30^ due to the post-mitotic nature of neurons, which makes them unamenable to clonal expansion and a source of extremely low frequency—often private to individual cells—mutations. These studies revealed that somatic mutations occur at higher rates in neuronal regulatory elements^26^, vary in rates and patterns between neurons and oligodendrocytes^21^, and occur at elevated rates in Alzheimer’s disease^27^ and multiple sclerosis lesions^30^. However, scWGA requires intact, high-quality, single nuclei, and protocols to isolate these nuclei depend on the tissue under study and the state of the cells. As such, applications of PTA to other tissues focused on cultured cells or cells from fresh specimens^25,31^; protocols for other postmortem tissues are under active development^32^. The SMaHT (Somatic Mosaicism across Human Tissues) Network is a multi-institutional collaboration to characterize somatic mutations across normal human tissues in postmortem donors. The Network will generate a wide variety of data types, including deep short-and long-read sequencing of bulk tissues, duplex sequencing of bulk tissues using several approaches, and scWGA^33^.

In this work, we describe a network-wide effort to isolate and sequence 102 single nuclei from frozen, postmortem lung and colon tissue of a 74-year old individual. A vast data resource was also generated by the Network for the same individual, which provided a rich catalog of orthogonal data to validate the accuracy of scWGA. PTA scWGA proved to be high-quality across multiple SMaHT-affiliated sites and tissues. Most importantly, single-cell resolution revealed unique insights that would normally require multiple experimental approaches and were missed by bulk tissue analyses (***The Somatic Mosaicism across Human Tissues Network et al., in prep #1***).

## Results

### Consistent, high-quality amplification of single nuclei across centers

Frozen tissue cores or homogenates from the lung and colon of a deceased 74-year-old male were distributed to four experimental laboratories in SMaHT (**Figure 1A**). Each laboratory developed protocols to isolate high-quality single nuclei from each tissue and to perform whole genome amplification by PTA (**Methods**). In total, PTA libraries for 102 cells (56 single lung cells and 46 single colon cells) passed initial quality control by PCR of four chosen loci (**Methods**) and were sequenced on the Illumina NovaSeq X platform at five sequencing centers, resulting in five datasets (**Table S1**). Sequencing data were collected at a single center for alignment, additional quality control, and somatic mutation calling. Sequencing depths ranged from 15×-50× per cell and 86% of the accessible genome was covered by ≥10 reads on average (**Figures 1B and 1C**). Post-sequencing quality control metrics designed to identify aberrant or failed amplification excluded 15 of the 102 PTA libraries (**Figures 1D and 1E; Figure S1; Table S2**), and all four laboratories were represented among the 87 retained cells.

**Figure 1.**
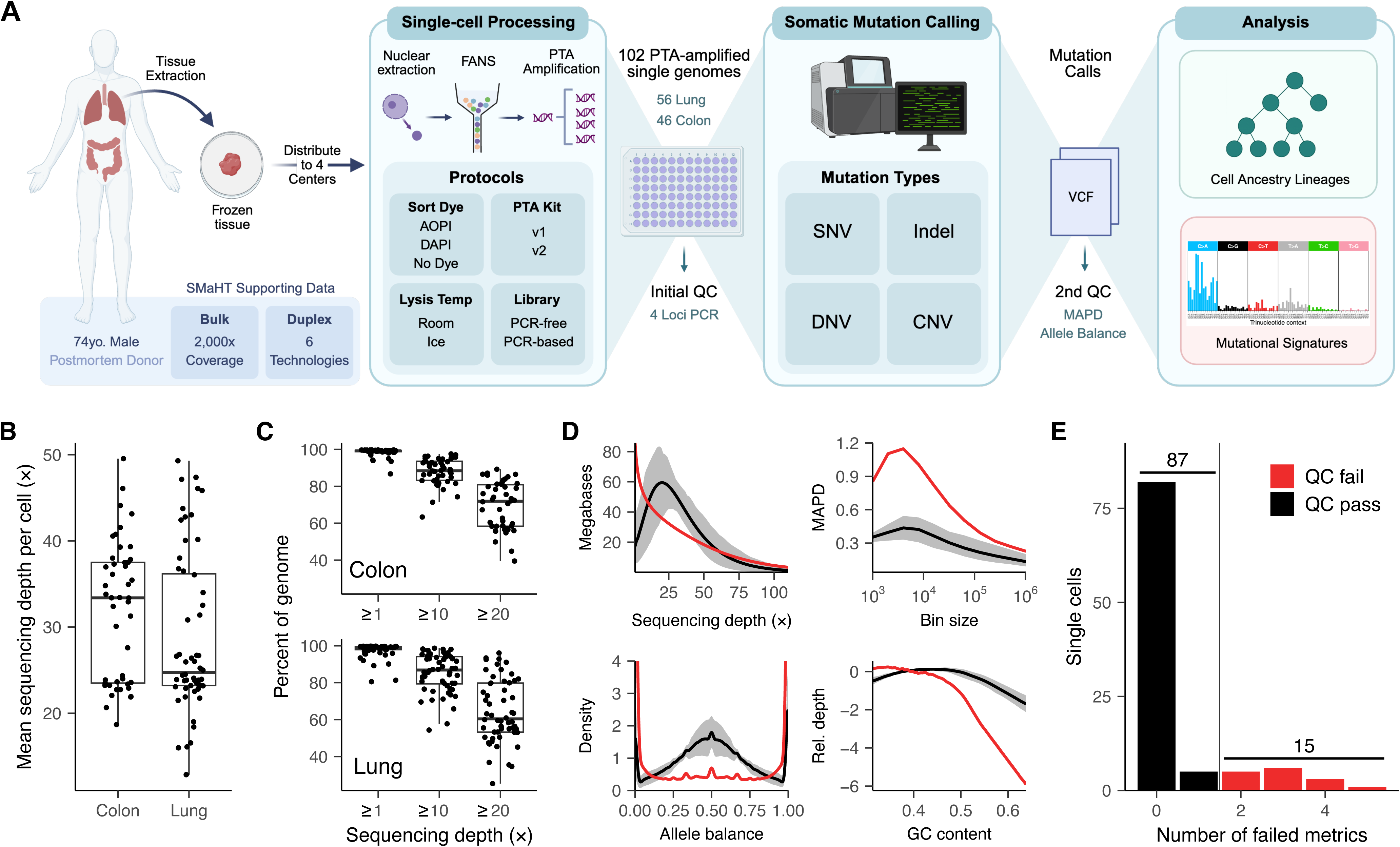
Experimental overview and quality control of PTA single-cell DNA sequencing. **A.** Experimental overview. Protocols for tissue collection, nuclei isolation, single-cell whole genome amplification and sequencing varied across centers. Final data were collected at a single center for uniform bioinformatic processing, secondary quality control, somatic mutation calling and analysis. **B.** Mean whole-genome sequencing depth per cell. **C.** Breadth of genomic coverage quantified by the percent of genome covered at various sequencing depths per cell. The total accessible genome was defined as the subset of GRCh38 with >0 coverage in Illumina sequencing of bulk from the same two tissues. **D.** Four examples of post-sequencing secondary quality control metrics. In each case, a rejected cell is depicted in red. For visual comparison, the average signal across all 102 cells is shown as a black line and the 10th-90th percentile range is shown as a grey envelope. **E.** Cells failing two or more secondary quality control metrics were excluded from SNV, indel and DNV calling and downstream analyses.

### Single-cell sequencing yields a high-quality somatic mutation catalog

Small somatic mutations—single nucleotide variants (SNVs), insertions and deletions (indels) and dinucleotide variants (DNVs)—were called using SCAN2^26^, which accounts for amplification-induced allelic imbalance to increase specificity of variant calling in PTA data. Deep (∼100×) Illumina bulk sequencing of lung and colon tissue of the same individual was used to remove germline variants. The vast majority of the genome satisfied the sequencing depth requirements (**Methods**) for SNV/DNV (mean 94% across cells) and indel (mean 88%) calling (**Figure 2A**). On average, SCAN2 recovered 1,385 SNVs, 76 indels and 17 DNVs per lung cell and 863 SNVs, 45 indels and 7 DNVs per colon cell (**Figure 2B**) resulting in mutational catalogs of 68,083 and 37,563 small mutation calls in lung and colon, respectively (**Figure 2C)**. Sensitivity for somatic SNVs and indels averaged 37% and 25%; and 2% of lung and 4% of colon somatic SNVs and indels were expected to be false positives based on previous benchmarking of SCAN2 performance (**Figure S2, Methods**). These low false discovery rates are reasonable given the relatively high mutation burden of cells from an individual of this age (74 y.o.).

**Figure 2.**
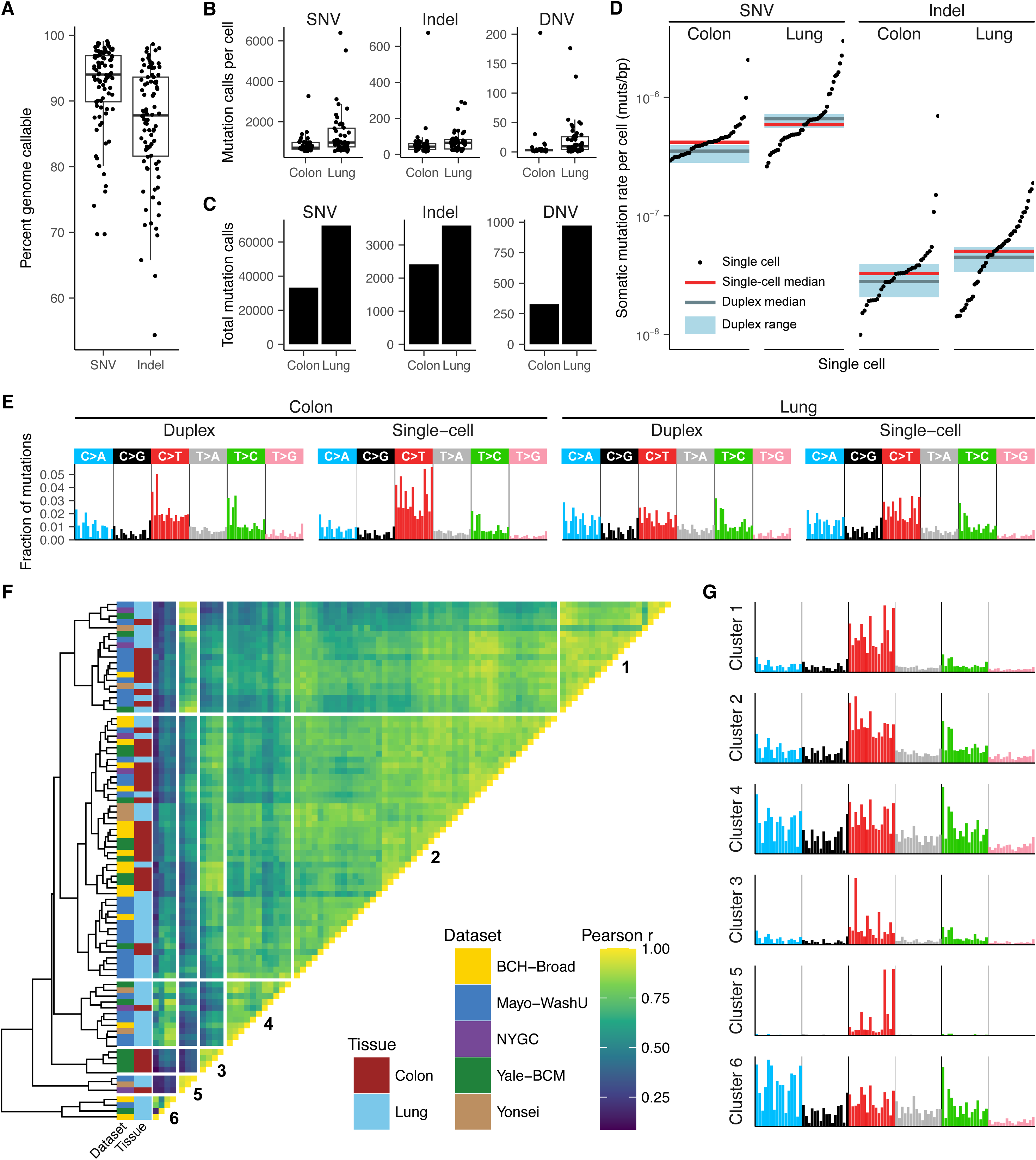
Single-cell small mutations recover mutation rates and spectra and reveal cellular heterogeneity. **A.** Percent of each single-cell genome with sufficient sequencing depth for small mutation calling by SCAN2. Sequencing depth requirements are identical for SNVs and dinucleotide variants (DNVs). Each point represents one single cell. **B.** Number of somatic mutation calls per single cell. **C.** Total number of somatic mutation calls for each tissue and mutation class. **D.** Somatic mutation rate per basepair, after correction for calling sensitivity, for each single cell. Horizontal lines indicate the median mutation rate across PTA single cells (red) and 6 duplex sequencing technologies (gray). Pale blue boxes indicate the minimum and maximum across the 6 duplex sequencing technologies. **E.** Comparison of combined 96-trinucleotide SNV mutational spectra between single-cell and duplex technologies. Spectra show fractions of mutations per channel. **F.** Hierarchical cell clustering based on Pearson’s correlation coefficient between SNV mutational spectra. The cluster dendrogram was cut at k=6 clusters. **G.** Average SNV spectrum across cells in each cluster identified in **F**.

To validate the accuracy of our single-cell mutation catalogs, we compared single-cell mutational burdens and spectra to those obtained by duplex sequencing of the same tissues. As part of a separate SMaHT effort, six duplex sequencing technologies, which correct artifacts by sequencing both DNA strands independently to detect mutations from single DNA molecules, were applied to homogenates of the same lung and colon specimens (***Zhang et al., in prep #3***). Average per-cell somatic SNV and indel mutation rates (corrected for calling sensitivity) and spectra using SCAN2 analysis were compared against those derived from duplex sequencing. Single-cell rate estimates were remarkably consistent with duplex-based measurements: the median single-cell SNV rates were 8.4% lower in lung and 16% higher in colon than the median of the 6 duplex technologies from the same tissue, whereas median single-cell indel rates were 13% and 10% higher than the corresponding median duplex rates for lung and colon, respectively (**Figure 2D**). The differences in median rates between single-cell and duplex technologies was much smaller than the maximum difference among duplex technologies themselves, which ranged from 130% (SNVs in lung) to 190% (indels in colon). Aggregate single-cell SNV mutational spectra for each tissue were also concordant with the aggregate spectra across duplex technologies, with cosine similarities of 0.97 and 0.91 for lung and colon, respectively (**Figure 2E**). Indel spectra were not compared since not all duplex technologies provide indel calls.

To investigate cell-to-cell mutational heterogeneity, we clustered single cells by SNV mutational spectra, defining at least 6 distinct clusters (**Figure 2F**). Clusters 1 and 2 were roughly evenly constituted by lung and colon cells, with the two clusters being differentiated by rates of T>C mutations (**Figure 2G**). The remaining four clusters were more tissue specific. Cluster 3 contained only colon cells and was distinguished by high rates of C>T substitutions at CpG dinucleotides. Clusters 4, 5 and 6 were dominated by lung cells. Clusters 4 and 6 exhibited widespread C>A mutations, with cluster 6 showing the most pronounced C>A burden and containing the three highest-burden lung cells. Cluster 5 was almost entirely defined by C>T mutations at the T<u>C</u>N trinucleotide context. The heterogeneity in mutation spectra amongst the cells indicated different active mutational processes, potentially reflecting cell type or cell history.

### Per-cell DNA damage process heterogeneity revealed by mutational signatures

Mutational signature analysis, which draws from curated databases of mutational patterns linked to mutagenic processes^34,35^, provided biological interpretations for some of the heterogeneity observed in our single cells. Six mutational components were extracted *de novo* using a composite approach (**Methods**, **Figure S3**) which combines analysis of three small mutation types (SNVs with 96 dimensions, indels with 83 dimensions and DNVs with 78 dimensions). The majority of mutations across cells was attributed to a *de novo* component 1, with a SNV pattern similar to COSMIC SBS5, a clock-like signature (**Figure 3A**). Component 4 captured COSMIC SBS1, another ubiquitous clock-like signature, and component 2 accounted for T>C mutations similar to COSMIC signatures SBS5 and SBS16. Lung cells had generally higher levels of the SBS5/SBS16-like component 2, while colon cells showed greater levels of the SBS1-like component 4, consistent with previous studies of lung^7^ and colonic epithelial cells^4^.

**Figure 3.**
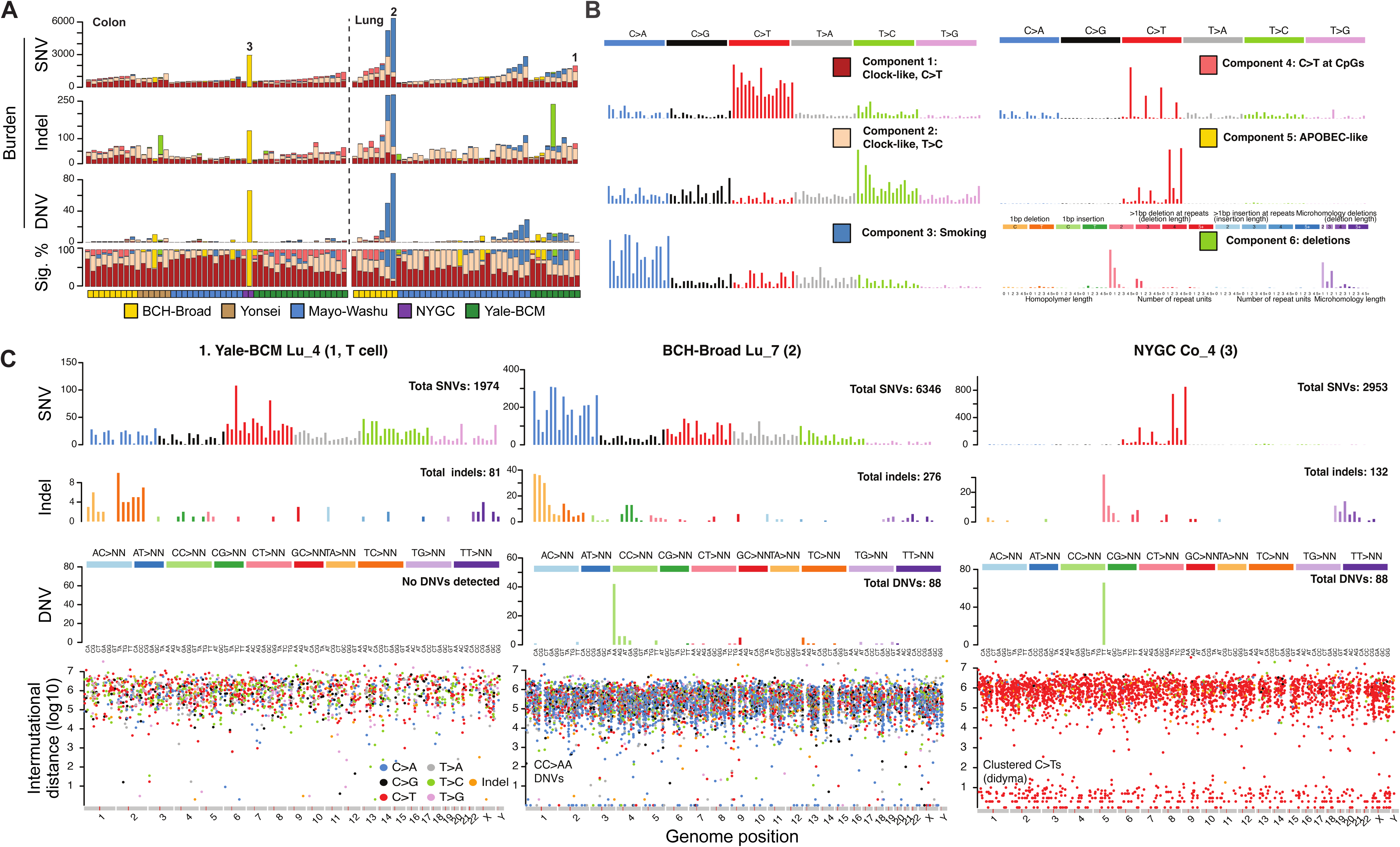
Signatures of tobacco smoking-and APOBEC-related mutagenesis at single-cell resolution. **A.** Exposures to each of the 6 *de novo*-extracted composite mutational components per single cell, annotated by tissue type and data set. The three numbered cells correspond to the cells depicted in panel (**C**). **B.** De novo-extracted composite mutational components. Composite components represent all three small mutation types (SNVs, indels and DNVs) encoded in a single 257-dimensional spectrum. **C.** SNV spectra and intermutational distance plots for 3 selected cells. (Left) T-cell from lung, (middle) lung cell with apparent tobacco smoking damage, and (right) colon cell with APOBEC damage. Numbering corresponds to panel (**A**). Rainfall plots (bottom row) show the distance between successive pairs of somatic mutations.

Two more components corresponded to signatures linked to specific mutagenic processes, suggesting unique mutational histories among our sampled cells. Component 3 was present primarily in lung cells (**Figure 3A**) and resembled COSMIC signatures spanning several mutation types caused by tobacco smoking: the C>A rich SBS4 SNV signature, the 1 bp deletion signature ID3 and the CC>AA dinucleotide DBS signature (**Figures 3B and 3C**), suggesting a single mutagenic etiology. Two lung cells showed exceptionally high component 3 mutation burden. These observations suggest a history of smoking (donor information is available at dbGaP), and presence in some but not all lung cells is consistent with previous observations in the lungs of both smokers and ex-smokers^7^.

Component 5 was identified in a handful of colon and lung cells and resembled SBS2, the C>T component of APOBEC-induced DNA damage. No kataegis—clusters of mutations created in the vicinity of double strand breaks—was observed in these cells; instead, pairs of C>T mutations separated by <100 bp were scattered across the genome (**Figures 3C** and **S4**). This is consistent with a recently observed^36^ interplay between APOBEC and tobacco mutagens that leads to mutations in pairs with intermutational distance less than 32 base pairs, termed *didyma* (Greek for *twins*). Occasionally, the pairs of C>Ts occurred directly adjacent to one another, creating a CC>TT dinucleotide variant and giving the false impression of exposure to the ultraviolet light-associated COSMIC signature DBS1, which is unlikely to be active in these cells. A single lung cell had both C>G and C>T APOBEC-driven mutagenesis but no evidence of *didyma* (**Figure S4**), suggesting differential activation of APOBEC3A and APOBEC3B across cells.

### Shared mutations from single cells reconstruct phylogenies to the zygote

Somatic mutations shared between different cells inform on the ancestry of these cells and can be used to reconstruct a phylogenetic tree. Mutations shared across a large number of cells, particularly when cells are from different tissues, represent early embryonic events that are likely detectable in bulk sequencing. Thus, the typical approach of removing candidate somatic mutation calls present in a matched bulk dataset would filter out early mutations. We addressed this by relaxing SCAN2’s filters against the matched bulk sample followed by the Sequoia phylogeny reconstruction algorithm^37^, which includes filters to remove recurrent artefactual mutation calls (**Methods**). These SCAN2 calls produced by relaxed bulk filters were only used for phylogeny analysis.

We identified 197 SNVs and 2 indels present in at least two cells and organized the single cells into a phylogenetic tree that reflected the early development of the donor (**Figure 4A**). Two lines of evidence support the accuracy of the single-cell derived phylogeny. First, cells from different datasets did not cluster, suggesting little or no bias between the various protocols for cell isolation, amplification and sequencing. Second, phylogenetic reconstruction using mutation calls from deep bulk whole genome sequencing of the same colon and lung tissues (***Jang et al., in prep #12***) produced a nearly identical topology (**Figure S5**). However, mutation discovery in bulk data missed a crucial early mutation, resulting in the omission of one of the earliest branching events. Overall, single-cell PTA captured most of the shared mutations discovered by bulk tissue sequencing (**Figure 4B**), establishing PTA as a bulk-independent approach for building cell phylogenies on par with LCM and cell cloning approaches^3,14,15,19,38^.

**Figure 4.**
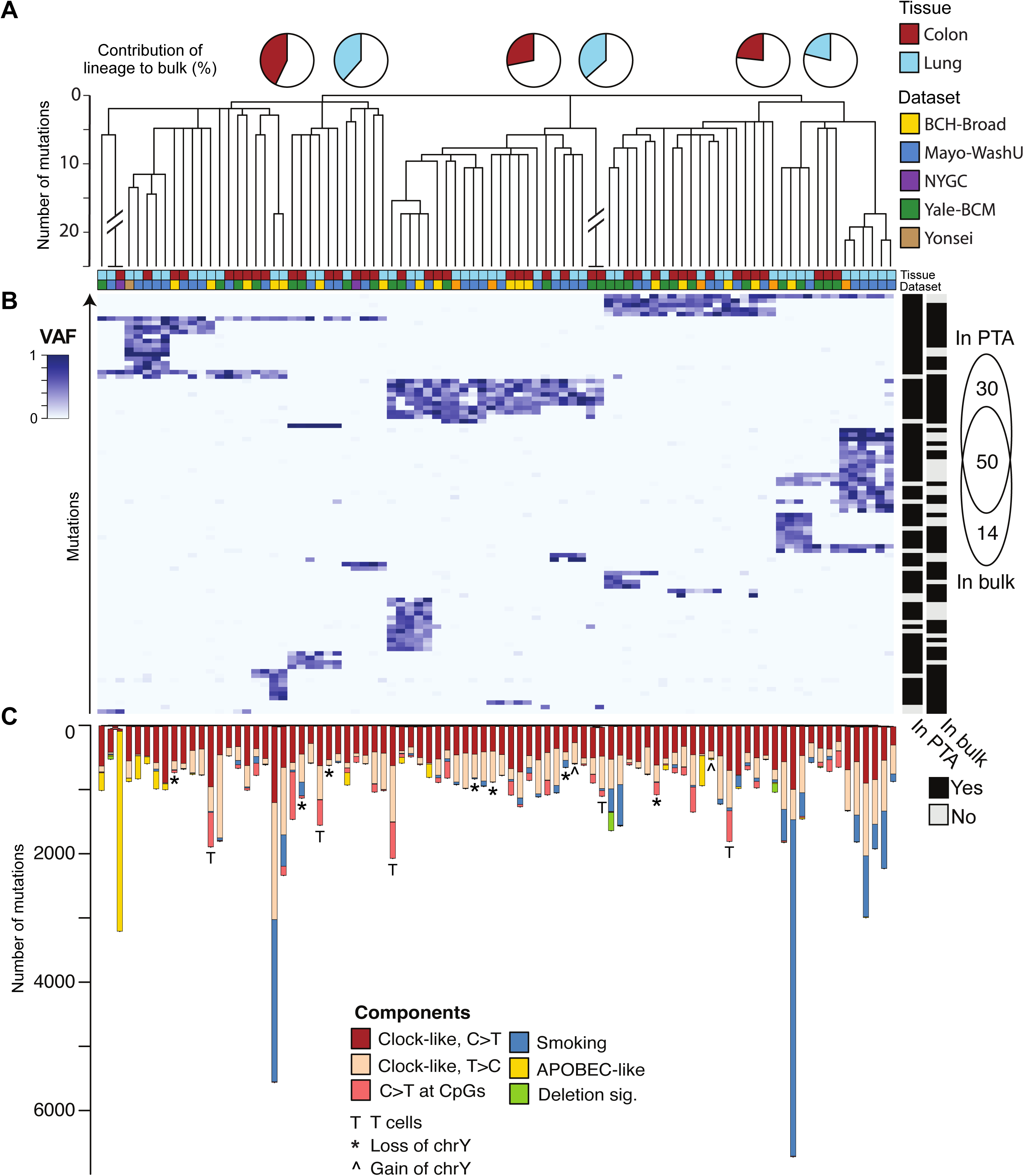
Cell lineages reconstructed by single-cell somatic mutations are confirmed by deep bulk sequencing. **A.** Phylogeny constructed from shared somatic mutations (SNVs, indels, DNVs) found in single cells reproduces cellular lineages rooted at the zygote. Pie charts show the share of the cell population represented by the three initial clades. Branch lengths are cut off at 25 mutations to emphasize the early phylogeny. **B.** Mutation VAFs in single cells at lineage informative loci. Cell order corresponds to panel (**A**). Two orthogonal approaches to identify shared/mosaic mutations are shown: one using only mutations called single cells (denoted as “In PTA”) and another using only mosaic mutation calls from deep bulk sequencing (denoted as “In bulk”). Each mutation is marked by which approach detected the site. A Venn diagram (right) demonstrates the overlap between the two approaches. **C.** Same phylogeny in (**A**) without a branch length limit. Branches are annotated with mutational component exposures defined in **Figure 3** and **Figure S3** and cells are annotated with aneuploidies of chromosome Y and re-arrangements at T-cell receptor loci, see **Figure 5**.

For both lung and colon cells, the most recent common ancestor was the root of the tree, assumed to be the zygote, which branched into three clades. By interrogating the VAF of mutations on the first three clades in bulk tissue sequencing data, we estimated the cell fraction for each clade-lineage. The combined cell fractions across the three lineages summed to approximately 100% in both lung and colon, indicating that the root of the tree likely represents the zygote. The broad topology of the tree matched those reconstructed previously in different individuals using single cell cloning and selection of highly clonal tissue specimens^14,15,19^. Branching of the root into three clades was also observed previously^15,39^, and suggests that either no mutations occurred in one of two daughter cells of the zygote or such mutations remain undetected, leaving the first branching unresolved. Finally, mutation burden and signatures showed no correlation with branches, with the clock-like SBS5 signature ubiquitously present in all cells, while exposure-induced signatures (i.e., smoking and APOBEC) occurred in adult life and were present in late branches of the phylogeny and without clade-level structure (**Figure 4C**).

### Frequent genomic rearrangements and aneuploidies in single cells

We performed copy number alteration (CNA) calling using CNVpytor^40^ and HiScanner^41^ in all cells, regardless of the quality control filters used for SNV and indel analysis. CNVpytor and HiScanner detect haplotype imbalances by leveraging read depth (RD) and B-allele frequency (BAF), classifying events as deletions (DELs), duplications (DUPs), or copy number neutral loss of heterozygosity (LOH). Most cells exhibited only a few CNAs—up to 6 events—with less than 15% of the genome affected (**Figure S10**). However, 4 cells presented with a substantially higher number of CNAs or a larger proportion of the genome altered. Visual inspection revealed extensive deletions at the chromosome or sub-chromosomal level (**Figure S6**), nearly all of which lacked breakpoint-spanning discordant reads. We thus suspected these cells experienced widespread amplification-associated dropout rather than *bona fide* deletions and focused our CNA analysis on the remaining 98 cells.

Among these 98 cells, 14 exhibited full-chromosome aneuploidies, with the most common event being loss (6 cells) or gain (2 cells) of chromosome Y (**Figures 5A** and **S7**). These events occurred in cells from lineages that diverged early in development, indicating that each gain or loss event arose independently (**Figure 4C**). This aligns with known biology: loss of chromosome Y is the most common somatic aneuploidy in males^42^. Interestingly, 6 of the 14 aneuploid cells showed copy number changes involving multiple chromosomes (**Figure S8**).

**Figure 5.**
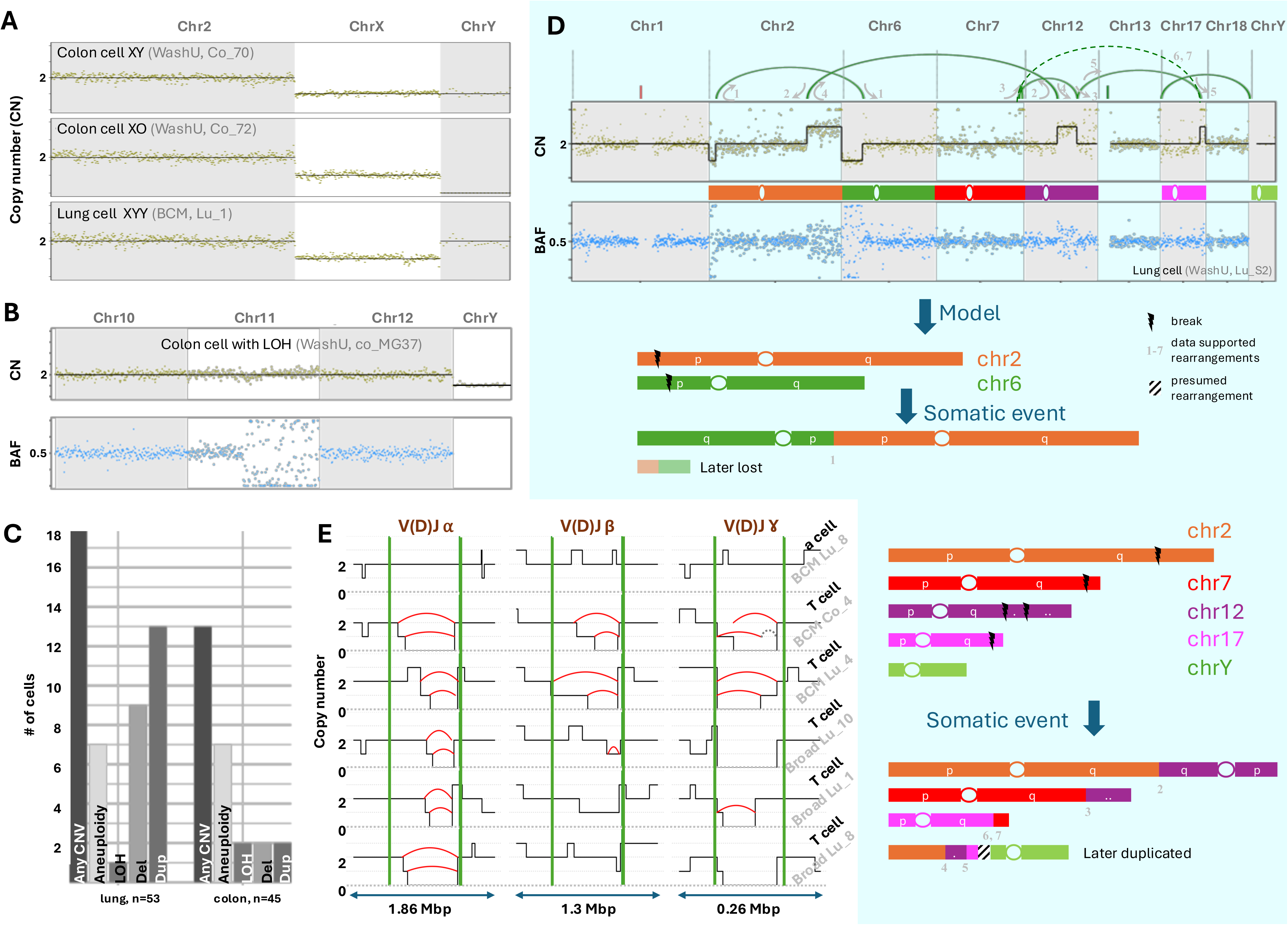
Copy number alterations (CNAs) in single cells. **B.** Read depth profiles for 3 representative cells including 2 with chromosome Y aneuploidy, which was the most common aneuploidy observed. Chromosome 2 was selected to represent autosomes. **C.** Read depth and B-allele frequency (BAF) plots for a cell with copy-neutral loss of heterozygosity. Large, multi-megabase-scale loss of heterozygosity (LOH) events are unlikely to be caused by random amplification biases. **D.** Number of cells with aneuploidies or subchromosomal CNAs >10 Mb in size. Number of cells listed is after excluding four cells following CNA QC. **E.** (Top) Read depth and BAF plots for a single cell harboring multiple CNAs supported by breakpoint junction-spanning reads. Chromosomal rearrangements inferred from discordant and split reads are shown by arcs (red for deletions and green for other rearrangements). Rearrangement directions are shown by arrows. (Bottom) A model of chromosome rearrangements to explain the observed data. Several translocations form non-canonical and dicentric chromosomes. Fragments from multiple chromosomes are likely fused with chromosome Y for a parsimonious explanation of their copy number gain, though no reads spanning the chromosome 17-Y junction were observed. **F.** Rearrangements in T-cell receptor loci (boundaries indicated by green lines) in five cells, with inferred copy number shown with black lines. Rearrangements were called from analysis of discordant and split reads (red arcs). In a few cells rearrangements were not called but are apparent from zero coverage. Split reads supporting an additional rearrangement were manually identified (gray dotted arc) to explain a rearrangement ending in a homozygous deletion.

We next analyzed subchromosomal CNAs ≥10 Mb in size. These CNAs were detected in 31 cells (13 in colon, 18 in lung), including several cells that also exhibited whole-chromosome aneuploidies (**Figures 5B-D** and **S8**). We observed three copy-neutral LOH events, which are likely real (**Figure 5B**), since a similarly-appearing artifact would require the unlikely scenario of complete dropout of one haplotype alongside exact and uniform doubling by overamplification of the other across megabases, far beyond the typical ∼2 kb PTA fragment size^25^. Similarly, DUP calls are likely genuine due to the observed BAF of 1/3 and 2/3; if they were artifacts, reproducing the expected BAF would require precise doubling by overamplification of one haplotype while maintaining normal amplification of the other. Large (>10 Mb) DELs likely reflect true biological events for the same reason: BAFs nearly 0 or 1 were observed over very large genomic regions.

Analysis of chromosomal rearrangements in the lung cell Mayo-WashU Lu_S2 provided orthogonal evidence confirming the accuracy of our read depth and BAF signals (**Figure 5D**). To resolve breakpoints of the 3 DUPs and 2 DELs (some smaller than 10 Mb) observed in this cell, we further analyzed it using Manta^43^, which searches for reads that span breakpoint junctions and offers orthogonal confirmation for RD and BAF analyses. It has been shown previously that both germline and somatic CNAs supported by RD/BAF and discordant/split reads are genuine genomic events^22,44^. The discordant and split reads recovered by Manta revealed that none of the CNAs were simple DELs or DUPs but rather represented complex chromosomal rearrangements (**Figures 5D and S9**). A model of genome alterations to explain the SV calls suggested several translocations leading to formation of non-canonical dicentric chromosomes that have been observed in various cancerous and non-cancerous samples^45^. The model explained all rearrangements and hypothesized only one fusion (chromosome Y with fragments from other chromosomes) not directly supported by junction-spanning reads. This fusion enables a parsimonious explanation for the extra copy of each of the chromosomal fragments and chromosome Y. Our model also posits molecules with telomeres on both ends, which is crucial for genome stability. The consistency of the rearrangement model with known biology supports the veracity of PTA-derived CNAs <10 Mb in size and highlights the unique perspective accessible through single-cell approaches by finding a cell with cancer-like genomic rearrangements in nominally healthy tissues^46^.

Finally, analysis of smaller SVs revealed 5 likely T lymphocytes among the cells isolated from lung tissue. We first searched for simple DELs and DUPs >100 kb using discordant and split reads and then further refined these candidates through RD and BAF analysis and inspection (**Methods**). This analysis revealed 22 deletions, 16 of which occurred at T-cell receptor loci (TCRs) in five cells (**Figures 5E and S10**). We hypothesize that the observed deletions occurred in T-cells as a result of somatic V(D)J recombination. Indeed, RD/BAF analysis confirmed that each of these cells has heterozygous and homozygous deletions in at least two TCRs indicating overlapping but different rearrangements on the paternal and maternal haplotypes. Furthermore, heterozygous deletions in three TCRs (alpha, beta, and gamma) were present in 4 cells. Taken together, these findings indicate somatic V(D)J recombination occurred in these cells, strongly suggesting that they are lymphocyte T-cells. The remaining two DUPs and four DELs outside of the TCRs highlight the ability to detect sub-megabase scale CNAs in PTA data (**Figure S11**).

## Discussion

We successfully applied PTA to normal cells from post-mortem lung and colon, while other studies have demonstrated PTA’s effectiveness in postmortem brain^21,26,27^, fresh blood^31^, and skin fibroblasts (***Natu et al., in prep #17***). Analysis of mutational signatures in single cells detected heterogeneous DNA damage profiles, highlighting the advantage of single-cell resolution, while also revealing expected tissue-specific signatures. Our study included PTA and duplex sequencing of the same tissues, allowing direct comparison. Duplex sequencing, an orthogonal approach that measures average characteristics of somatic SNVs and indels across cells, validated the mutation burdens and spectra from PTA, indicating that applied analytical methods designed to recognize PTA amplification artifacts are effective. However, duplex sequencing only partially captured heterogeneity among cells (e.g., tobacco exposure), underscoring the unique insights accessible with single-cell resolution.

Mutations derived exclusively from single-cell data also recovered cellular phylogenies with similar resolution to laser capture microdissection (LCM) of small clones from tissue and *ex vivo* clonal expansions of single cells. In previous efforts to study cell lineage relationships using single-cell data, expensive deep bulk sequencing played a central role in determining lineage-informative loci^5,47^. Our PTA-only single-cell phylogeny was validated by bulk sequencing and captured all major lineages after the first zygotic division. We observed a nearly symmetric contribution of early developmental lineages to tissues, which was observed in previous studies^14,15,19^. The accuracy of our phylogenetic tree suggests that allelic dropout during single-cell amplification—in which evidence of mutation sharing can be randomly lost—is not a significant concern when analyzing many cells.

A remarkable feature of PTA single-cell DNA sequencing is the ability to detect diverse mutation types, demonstrated through our analysis of large somatic mutations like CNAs and SVs. From PTA data alone, we reconstructed complex chromosomal rearrangements using two orthogonal forms of evidence:

*(1)* sequencing depth and (*2*) DNA fragments covering the junctions of rearrangements in individual cells (i.e., “split” or “discordant” reads). It is difficult for other amplification techniques to recover both of these signals, and LCM and *ex vivo* cloning may select against cells with extensive rearrangements if their proliferative potential is reduced. Further, since the CNAs and SVs we detected were not shared among our 102 cells, their rarity among the cell population likely excludes their detection using bulk-based technologies. Future work could leverage PTA’s simultaneous detection of SNVs and indels to explore the times at which large mutations arise by cross-referencing them with phylogenetically resolved small mutations on the altered large alleles.

Several forms of genetic heterogeneity captured by PTA enabled instances in which cell histories or cell type could be inferred. For example, mutational signature analysis revealed cell histories representing tobacco smoking and canonical APOBEC mutagenesis; inter-mutational distances indicated yet another cell history in which tobacco carcinogens and APOBEC had jointly produced pairs of C>T mutations through single events. CNA and SV analysis identified specific cell types by revealing rearrangements of the T-cell receptor locus in a handful of cells, which, when sampling heterogeneous cell types, as in the case of every tissue in the human genome, is one of the key pieces of information for understanding mutation heterogeneity across cells. Other signals derived from somatic mutations may further enable cell type inference: for example, the cell type of the ancestral founder of tumors was inferred by correlating the density of somatic mutations with cell type-specific histone modification and chromatin accessibility patterns^48^.

In summary, our study establishes PTA as a single-cell genomics platform applicable to post-mortem and fresh tissues, diverse cell types, and both proliferating and terminally differentiated cells to comprehensively survey mutation classes across the human lifespan. These properties position PTA as a universal technique for studying somatic mosaicism. Despite many capabilities of single-cell DNA sequencing, its current limitation is throughput and cost. An important complementary approach is duplex sequencing, which benefits from simpler bench protocols and subsequent data analysis. When applied to bulk it cannot generally resolve mutations to individual cells, challenging phylogeny reconstruction and possibly prohibiting detection of mutations that cannot be captured on a single molecule such as large CNVs. However, duplex sequencing excels in SNVs and indel mutational signature analysis and mutation rate estimation. We suspect future work to benefit from both single-cell and duplex methodologies, with PTA-based single-cell approaches casting a wider net and acting as a comprehensive toolkit for investigating somatic mosaicism across human tissues.

## Data Availability

The datasets described in this study are available through dbGaP under the study accession phs004193. The data used in this work were provided by the SMaHT Data Analysis Center (DAC) on behalf of the SMaHT Network. More information about the SMaHT Network and data is available online at https://smaht.org and at https://data.smaht.org.

## Code Availability

Scripts to perform analyses in this manuscript are available at doi:10.5281/zenodo.17413251. All bioinformatic tools are freely available for academic use.

## Acknowledgements

We are extremely grateful to the SMaHT donors, and donor families, who have generously provided such precious gifts to support this important work. This research is supported by the NIH Common Fund, through the Office of Strategic Coordination/Office of the NIH Director under awards U24 MH133204, U24 NS132103, UG3 NS132024, UG3 NS132061, UG3 NS132084, UG3 NS132105, UG3 NS132127, UG3 NS132128, UG3 NS132132, UG3 NS132134, UG3 NS132135, UG3 NS132136, UG3 NS132138, UG3 NS132139, UG3 NS132144, UG3 NS132146, UM1 DA058219, UM1 DA058220, UM1 DA058229, UM1 DA058230, UM1 DA058235, and UM1 DA058236.

## Methods

### Single-nuclei isolation and amplification by primary template-directed amplification scWGA at Boston Children’s Hospital

Approximately 10 mg of frozen post-mortem tissue homogenate for lung and colon were resuspended in homogenization buffer (10mM Tris-HCl pH 8, 250mM sucrose, 25mM KCl, 5mM MgCl2, 0.1% Triton X-100, Roche EDTA-free protease inhibitor), Dounce homogenized, filtered through a 40 um mesh filter, and centrifuged in a swinging-bucket centrifuge for 10 minutes at 900 rcf. Nuclei were resuspended in PBS with 0.8% BSA and incubated with DAPI (final concentration: 100 ng/mL). Single DAPI+ nuclei were isolated using fluorescence-activated nuclei sorting on a BD FACSymphony S6. Whole-genome amplification was performed using the ResolveDNA™ PTA V1 kit V1 (BioSkryb).

### scWGA at Yale University

∼10mg of tissue was chopped into small pieces and resuspended in lysis buffer (319.6mM sucrose, 5mM CaCl2, 3mM Mg(Ac)2, 0.1mM EDTA, 10mM Tris-Cl pH 7.4, 1mM DTT, 0.1% TritonX-100). Nuclei were extracted by douncing tissue pieces in lysis buffer. Suspension was filtered through 40 m filter, and the flow-through was centrifuged at 500 rpm for 10 mins at 4 °C. The pellet was resuspended in sorting buffer (1% BSA, 1 mM EDTA, 10 mM HEPES prepared in 1X PBS) and stored on ice until sorting. Extracted nuclei were subjected to sorting through 100 m Nozzle and without any fluorescent staining. Nuclei from a human fibroblast cell line was used for gating, followed by FSC-A/SSC-A gating for nuclear integrity and FSC-A/FSC-H for doublet exclusion. Nuclei from tissue samples were sorted based on the same gating into BioSkryb cell buffer. Whole-genome amplification was carried out as per ResolveDNA™ PTA V1 kit V1 (BioSkryb) recommended protocol.

### scWGA at Yonsei University

#### Nuclei isolation from frozen Lung tissue

10–30mg of frozen human lung tissue was gradually thawed at 4 °C. Visible adipose tissue and blood vessels were removed, and the samples rinsed thoroughly with cold 1X PBS until no blood was visible. The tissue was minced into small fragments and transferred to a cartridge for nuclei extraction using Minute™ Detergent-Free Nuclei Isolation Kit (Invent Biotechnologies, Cat. No. NI-024). Tissue was lysed by grinding in pre-chilled buffer A followed by incubation on ice for 5–10 minutes for complete lysis. The nuclei-containing suspension was subjected to centrifugation at 16,000 × g for 20 seconds in a tabletop microcentrifuge. The resulting pellet was resuspended by vigorous vortexing for 10 seconds and centrifuged at 500 × g for 2–3 minutes. Pellet was washed with cold buffer B and centrifuged at 600 × g for 8–10 minutes. The resulting pellet was resuspended in ice-cold FACS buffer (PBS supplemented with 1% BSA) and processed for FACS-based single-nucleus sorting.

#### Nuclear isolation from frozen Colon tissue

Frozen human colon tissues were gradually thawed at 4 °C and washed with cold phosphate-buffered saline (PBS) until no visible blood remained. Under a dissection microscope, the mucosal layer—presumed to be enriched for colonic crypts—was carefully dissected using sterile fine-tipped forceps and transferred directly into the cartridge for nuclei isolation. The remaining tissue was cleared of adipose tissue and debris, placed onto a sterile glass plate, and finely minced using a sterile surgical blade until no further dissection was feasible. The inner layer, containing mostly crypt, was processed following the same nuclei isolation procedure used for lung tissue, based on the Minute™ Detergent-Free Nuclei Isolation Kit protocol. For outer layer, minced tissue fragments were transferred to a 1.5 mL microcentrifuge tube, and 100 μL of pre-chilled non-ionic detergent Cell Lysis Buffer II (Invitrogen, Cat. No. FNN0021) was added. Samples were incubated on ice for 10∼30 minutes and homogenized using a disposable pestle until the lysate became visibly clear. The suspension was centrifuged at 600 × g for 5 minutes at 4 °C, and the supernatant was discarded. The resulting pellet was gently resuspended in cold FACS buffer (filtered 1% BSA in PBS) and washed twice to remove residual lysis buffer and cellular debris. The final suspension was filtered through 40 μm cell strainer to reduce large debris. The final nuclear suspension was resuspended in cold FACS buffer for downstream applications.

#### Staining-based sorting of nuclei with dyes

To evaluate nuclear integrity and viability, samples were stained with either Acridine Orange/Propidium Iodide (AO/PI; Logos biosystem; Cat #F23011; 2ul AO/PI, 18ul sample) or SYTO9 (Thermo Fisher Scientific; 5 µM), following the respective manufacturer’s protocols. Staining was performed for 5–10 min at 4 °C in the dark. Nuclei were then gently washed in cold FACS buffer and passed through a 40 µm cell strainer (Falcon) to obtain a single-nucleus suspension. Flow cytometric sorting was conducted on a BD FACSAria III cell sorter (BD Biosciences) equipped with a 70 μm nozzle, operated at low pressure (∼20 psi) to reduce shear-induced damage. Nuclei were gated based on forward and side scatter (FSC-A and SSC-A), and doublets were excluded using FSC-H vs. FSC-A profiles. Fluorescent gating was performed using AO/PI or SYTO9 signal: PI events were selected in AO/PI-stained samples, and SYTO9 singlets were selected in SYTO9-stained samples. Fluorescence compensation was applied using single-stained and unstained controls. Sorted nuclei were collected into 96-well plates and immediately processed for PTA reaction.

#### Dye-free sorting of nuclei

In this approach, nuclei were not stained with any fluorescent dyes.

Instead, gating parameters were optimized using DAPI-stained control samples (5 µg ml ¹; Thermo Fisher Scientific) to define the forward and side scatter profiles (FSC-A vs. SSC-A) of intact nuclei and to exclude doublets and aggregates using FSC-A vs. FSC-H plots. These gating thresholds were subsequently applied to unstained nucleus samples during sorting.

#### scWGA at Weill Cornell Medicine

Tissue was loaded on Singulator along with final concentration of 1.0 U RNAse inhibitor (Protector, Sigma) and 1mM DTT using standard nuclei extraction protocol. Sucrose (final concentration 250 mM) was added to nuclei output to mix and spun down at 500 g, 4 °C, for 5 minutes. Nuclei were washed with a nuclei wash buffer (1X DPBS, 1.0 U RNAse inhibitor, 1mM DTT, 1% BSA) twice and filtered through a 40 μm strainer. Nuclei were stained with DAPI and counted on a Countess. https://www.protocols.io/view/nuclei-extraction-for-single-cell-rnaseq-from-froz-q26g74xzqgwz/v1

#### PTA reaction

Whole-genome amplification was performed using the ResolveDNA™ PTA kit V1 or V2 (BioSkryb) according to the manufacturer’s instructions in a DNA-free pre-PCR hood. Briefly, sorted 96 well plates were initiated by sequential addition of MS mix (1.5 µL SM2 + 1.5 µL 1× SS2), SN1, and SDX reagents with intermittent mixing at 1,400 rpm and room temperature incubation. The temperature during the lysis step with MS mix was RT or 4. For amplification, 8 µL reaction mix (SB4, SEZ1, SEZ2, 1× SS2) was added, and thermocycling was conducted at 30 °C for 10 h, 65 °C for 3 min, followed by a 4 °C hold. Amplified DNA was purified using ResolveDNA magnetic beads. Negative (buffer only) and positive (1–1,000 pg gDNA) controls were included in each run. All pipetting was performed with pre-chilled, low-retention filtered tips, and mixing was carried out using an RPM-controlled plate shaker.

#### Quantification of DNA

DNA concentrations were determined using the Qubit™ dsDNA High Sensitivity Assay Kit (Thermo Fisher Scientific, Cat. Nos. Q32851/Q32854) and Qubit™ Fluorometer, according to the manufacturer’s instructions.

#### Quality assessment of whole-genome amplified DNA

Quantity was determined using the Qubit HS dsDNA assay (Thermo Fisher), and the quality of genome amplification was assessed by the 4-loci test (at Yale University, Yonsei University, and Boston Children’s Hospital).

#### Primer design

To evaluate the quality and genomic representation of whole-genome amplified (WGA) DNA, a 4-locus multiplex PCR assay was developed. 4 loci were selected from chromosomes 5, 10, 15, and 20, yielding non-overlapping amplicons ranging from 200 to 500 bp (**Table S3**).

#### Sample selection

Samples were subjected to a four-locus quality control assay, and only those yielding all four expected amplicons were deemed to have sufficient genomic representation and retained for downstream analysis. These samples exhibited DNA concentrations exceeding 100 ng, as determined by Qubit™ dsDNA quantification.

#### Illumina sequencing at the Broad Institute

The PCR-Free whole genome sequencing (WGS) process includes sample preparation utilizing custom Broad indices, which are commercially available through Roche and Kapa Biosciences HyperPrep library construction kit, sequencing on the NVX 25B (Illumina 2×150bp reads), read de-multiplexing, aggregation, and alignment (DRAGEN).

#### Illumina sequencing at Washington University, St. Louis (WashU)

Amplified genomic DNA samples were quantified using the Qubit Fluorometer. Genomic DNA (∼250ng) was fragmented on the Covaris LE220 instrument targeting ∼375bp inserts. Fragmented DNA was size selected using 0.8X ratio of Ampure XP beads (Beckman Coulter) to remove fragments less than 300bp. Dual indexed libraries were constructed utilizing the KAPA Hyper library prep kit (Roche Diagnostics, Cat # 7962363001). Full length custom adaptors were used during ligation (IDT, UDI/UMI configuration with 10bp UDIs and a 9bp UMI in the i7 position) then amplified for 8 PCR cycles. Libraries were run with KAPA Library Quantification kit (Roche Diagnostics) to measure molar concentration. Libraries were sequenced on NovaSeq X using paired end reads extending 150bp targeting >30X coverage.

#### Illumina sequencing at Yonsei University

Sequencing libraries were constructed from 1 μg amplified nuclear DNA using the TruSeq DNA PCR-free library kit (Illumina, San Diego, USA) according to the manufacturer’s instructions. DNA was fragmented using the Covaris system (Covaris, Woburn, Massachusetts, USA) and sequenced to a target depth of 30x on a NovaSeq X plus platform (Illumina, San Diego, USA).

#### Illumina sequencing at New York Genome Center (NYGC)

Primary Template-directed Amplification (PTA) libraries were prepared using NEBNext Ultra II DNA PCR-free Library preparation Kit (NEB E7410L) in accordance with the manufacturer’s instructions. 250ng of PTA product was fragmented mechanically using the Covaris R230 to ∼350bp fragments. DNA fragments were subsequently end-repaired and adenylated. DNA fragments were ligated to Illumina sequencing adapters (Twist Bioscience) and the libraries underwent bead-based size selection. Final libraries were quantified using the QuantStudio5 Real-Time PCR System (Applied Biosystems) and Fragment Analyzer (Agilent).

#### Illumina sequencing at Baylor College of Medicine (BCM)

Following samples QC using Pico green DNA is aliquoted to 200ng in 70 ul and is sheared into fragments of approximately 200-600 bp in a Covaris E220 system. The shearing step is followed by double size selection and purification of the fragmented DNA using AMPure XP beads. DNA end repair and 3’-adenylation are performed in the same reaction followed by using automated processes with KAPA Hyper reagents (Roche# KK8505). A set of 96 Unique dual index barcodes (Illumina TruSeq UD Indexes, # 20040870) are utilized to barcode samples. Libraries were amplified for 8 PCR cycles in 50 μl reactions containing 150 pmol of P1.1 (5’-AATGATACGGCGACCACCGAGA) and P3 (5’-CAAGCAGAAGACGGCATACGAGA) primer and Kapa HiFi HotStart Library Amplification kit (Cat# kk2612, Roche Sequencing and Life Science). Fragment Analyzer (AgilentTechnologies, Inc) electrophoresis system is used for library quantification and size estimation. Prepared libraries are then pooled to achieve 30X coverage and sequenced on NovaseqX 25B flowcell (2×150bp reads).

#### Alignment and preprocessing of single-cell and bulk short reads

Illumina paired-end short reads were processed according to the Genome Analysis Toolkit (GATK) Best Practices. Prior to alignment, failed reads indicated by polyG repeats, thought to be caused by Illumina’s two-color chemistry, were filtered using fastp^49^ v0.23.2. The remaining reads were aligned to GRCh38 (GCA_000001405.15), which excludes alternate contigs and decoy sequences from hs38d1, using Sentieon’s^50^ (v202308.01) re-implementation of the BWA-MEM algorithm. Following alignment, PCR duplicates were marked by Sentieon *LocusCollector* and *Dedup*, local realignment around indels was performed by Sentieon *Realigner*, and base quality scores were recalibrated by Sentieon *QualCal* using dbSNP v138 and the Mills indel set—both provided in the GATK resource bundle—as databases of known polymorphic sites. Aligned and processed BAMs were then stripped of the indel quality score tags BI and BD to reduce file size, since they are disregarded during GATK HaplotypeCaller’s local reassembly.

#### Small somatic mutation detection and analysis with SCAN2

Small somatic mutations—single nucleotide variants (SNVs), insertions and deletions <50 bp (indels) and dinucleotide variants (DNVs)—were called using SCAN2 (http://github.com/parklab/SCAN2 commit f0c8f3f and http://github.com/parklab/r-scan2 commit da84a81), a somatic mutation caller developed specifically for analysis of PTA-amplified single cells. Full scripts to perform the analyses described below are provided in Zenodo. Common `scan2 config` arguments include: the same GRCh38 FASTA file used for alignment (`--ref`) and dbSNP v151 common (`--dbsnp`), `--gatk sentieon_joint` to use Sentieon’s implementation of GATK HaplotypeCaller (`driver --algo Haplotyper`) and `--genome hg38`.

#### Cross-sample panel construction

To enable somatic indel calling in SCAN2, a panel of bulk and single-cell data from several individuals is required. Since our data was generated from a single individual, 52 PTA single-cells and 18 matched bulks from an additional 17 individuals were downloaded from dbGaP (accession phs001485.v3.p1). To parallelize panel construction, a fine set of 100 kb bins spanning chromosomes 1-22, X and Y, excluding pseudo-autosomal regions and regions with anomalously high sequencing depth (totalling 8.3 Mb, see **Table S4**), was created. `scan2 makepanel` was then configured with this 100 kb bin set (`--regions-file`) and all BAMs from this study (102 single cells and 2 matched bulks) and the previous study (52 single cells and 18 matched bulks) supplied via the `–-bam` argument.

#### Mutation calling

Small mutations were called with `scan2 call_mutations` separately for colon and lung to allow single cells to be matched with bulks from the opposite tissue, reducing the likelihood that SCAN2’s filters would discard clonal mutations due to support in bulk. Importantly, SCAN2 determines read support for mutations by analyzing all BAMs simultaneously rather than the standard GVCF approach, in which BAMs are analyzed individually. The GVCF approach is much more computationally efficient but loses information by omitting read support for mutations if the support is low. Thus, to ensure identical read count matrices across analyses, all 104 BAMs were supplied to each tissue-specific analysis, but with different options specifying how each BAM was to be used. For example, to indicate that lung single cells should be genotyped relative to bulk data from colon, `--bulk-bam` was set to the bulk BAM from colon, the 56 lung single cell BAMs were supplied via `–-sc-bam` (one per BAM) and the lung bulk and 46 colon single cell BAMs were supplied via `–-bam` (one per BAM). The opposite assignment was used to call mutations in colon cells. `scan2 call_mutations` was additionally configured to use Eagle v2.4.1 for phasing (`--phaser eagle`) with the 1000 genomes reference panel, the 100 kb analysis bins used in panel construction (`--regions-file`) and the cross sample panel produced by `scan2 makepanel` (`--cross-sample-panel`). The individual’s sex was set to male (`--sex male`) and the amplification type for each cell specified by `--sc-bam` was set to PTA via `--amplification <cell ID> PTA`.

#### Mutation rates

Mutation rates are automatically estimated by SCAN2 during the `call_mutations` step. Rates were extracted from SCAN2 objects by first `load()`ing the object `obj` into R and running `mutburden(obj)/(get.gbp.by.genome(obj)*1e9)`.

#### Mutation calls

Mutation calls were extracted from SCAN2 objects in R by running `passing(obj, combine.mnv=TRUE)` after `load()`ing the SCAN2 object `obj` from disk. The `combine.mnv=TRUE` option enables DNV detection.

#### SNV and indel false discovery rate estimation

The rate of false positives was previously^26^ estimated to be 0.013 SNVs and 0.00073 indels per megabase of accessible genomic reference sequence using a synthetic data technique that preserved PTA amplification artifacts. The number of false positive calls per cell *FP*_mut_ is then the false positive rate multiplied by the number of bases analyzed by SCAN2 in that cell for mutation type mut. The final false discovery rate is FDR_mut_ *= FP*_mut_ / min(*FP*_mut_, *N*_mut_), where *N*_mut_ is the total number of mutation calls from the cell of type mut.

#### Sequencing data-driven quality control

The R objects output by SCAN2 allow for convenient interrogation of several quality metrics via the `scan2` R package and SCAN2’s summary objects, which summarize the full analysis objects and enable loading of 100s-1000s of cells for simultaneous analysis. In the commands that follow, `objs` is an R `list` of SCAN2 summary objects for all 102 single cells, loaded by SCAN2’s `load.summary()` function. MAPD, computed at several variably-sized bins that aim to capture equal numbers of alignable bases, was extracted using `mapd(objs)`. Allele balance, represented by the distribution of VAFs at heterozygous germline SNPs, was extracted with `ab.distn(objs)`. The read depth distribution, which is the number of sites in the genome with coverage = 1, 2,…is given by `dp.distn(objs)`. To calculate GC bias, the genome is tiled with non-overlapping 1 kb bins. The fraction of C and G nucleotides and the number of reads is computed per bin and a loess curve is fit to the number of reads per bin as a function of the GC content per bin. This Loess curve fit is returned by SCAN2’s `gc.bias(objs)`. Finally, read depth profiles along the genome were extracted using `binned.counts(objs, type=’cnv’)`. For each metric, all cells were plotted together and outlier cells were visually identified. Outlier status was then compiled across all 5 metrics and any cell with outlier status in 2 or more metrics was excluded from small mutation (SNV, indel and DNV) analyses.

#### Comparison to duplex-sequencing

Results from six duplex-sequencing technologies applied to the lung and colon tissue of our donor were available at the time of this study from ref. (***Zhang et al., in prep #3***): (1) NanoSeq using restriction enzymes (Hpy), (2) NanoSeq using mung bean nuclease (MBN), (3) CompDuplex using Tn5, (4) CODEC DRv1, (5) CODEC DRv2 and (6) CODEC ddBTP-Hpy. SNV and indel mutation rates were taken from the summary table for tissues, using the donor ID ST002 and tissue codes 1D (lung) and 1G (colon). Indel mutation rates for NanoSeq Hpy were excluded for being highly discordant with the other 5 technologies.

#### Single-cell lineage reconstruction by Sequoia

To avoid discarding early somatic mutations that define cell phylogenies near the zygote, it was necessary to relax SCAN2’s filters that require no read evidence of a somatic mutation in the matched bulk sample. Each single cell was relaxed by `load()`ing its SCAN2 object in R and running `update.static.filter.params()` with new parameters `max.bulk.alt=100`, `max.bulk.af=1`, `max.bulk.binom.prob=1e-5` for both SNVs and indels. The `alt` and `af` parameters were chosen to essentially disable those filters, shifting reliance primarily to the `binom.prob` filter. After running the update function, relaxed calls were extracted from the objects in the same way as non-relaxed calls (see *Small somatic mutation detection and analysis with SCAN2*) and a list of unique mutations, identified by chromosome, position, reference and mutated sequence, was identified across cells. The numbers of reference-and mutation-supporting reads were then retrieved for each cell and site from each cell’s SCAN2 object to create (site × cell) matrices of reference and mutation read counts. These matrices formed the input for Sequoia^37^, which further filtered the relaxed mutation calls and inferred the phylogenetic tree underpinning the single cell PTA data. To account for artificial overdispersion introduced by research site-specific variation, we expanded the beta-binomial overdispersion based filtering to calculate the maximum likelihood overdispersion parameter (‘rho’) across all cells, and within cells from each center. Alternative read count abundances for mutation calls shared between centers or broadly present within centers were required to be overdispersed (rho>0.3) between cells for each center they were present in.

#### Comparison of single-cell and bulk-derived cell lineage reconstruction

Somatic mutations were identified in bulk WGS of the same lung and colon tissues using a pipeline developed in ref. (***Jang et al., in prep #12***) for high depth, multi-tissue bulk samples. These loci were then interrogated in single-cells. First, a list of bulk derived SNVs was compiled and used to generate per cell pileups. At each locus, reads were required to meet a minimum mapping quality score of 20 (–q 20) and a base quality score of 20 (–Q 20) before counting reference and alternate alleles. The counts of reference-and alternate-supporting reads were assembled into a (variant × cell) matrix and fed into the Sequoia tree building pipeline, similar to the phylogenetic tree reconstruction for SCAN2-derived mutation calls, and mutations were mapped onto individual branches. Branches without any mutations were collapsed into polytomies. The comparison of phylogenies from bulk-or single cell-derived mutation calls was performed with the R package phytools. A normalized, generalized Robinson-Foulds distance was computed to compare the topology of the two trees using the JaccardRobinsonFoulds metric from the R package TreeDist (gRF=0.22).

#### Mutational signature analysis

The R package HDP (https://github.com/nicolaroberts/hdp), based on the hierarchical Dirichlet process, was used to extract mutational signatures. As a basis for mutational signature extraction, we used the counts of 257 categories across mutation types: 96 SNVs with trinucleotide contexts, 83 indels within their length and repeat/microhomology characteristics, and the 78 distinct types of DNVs. The mutational signature analysis was performed with mutations assigned to phylogenetic tree branches as distinct samples to avoid double counting of mutations. A hierarchical structure is established using sequencing centers as the first tier (parent nodes) and individual phylogenetic tree branches as the second tier (dependent nodes). If a tree branch led to cells from different sequencing centers, the parent node was set as ‘Shared’. In total, six distinct composite mutational signatures were extracted by HDP (**Figure S3**).

### Copy number alteration detection

#### Running CNVpytor

CNVpytor^40^ can utilize either read depth (RD) coverage, B-allele frequency (BAF) imbalance, or both signals for CNA analysis. First, genome segmentation is conducted using only the BAF signal. Segments with balanced BAF (i.e., BAF ∼0.5) are used to evaluate coverage for the normal diploid genome. In the next step normal segments are used to conduct GC correction across the genome. Then segmentation of the whole genome is performed. For the analysis of single-cell PTA data we utilized a 2-dimensional approach that integrates both RD and BAF signals during segmentation.

To calculate the BAF signal we obtained germline variants called from a bulk sample. SNPs in this donor were phased using shapeit4^51^. In each cell, BAF was calculated per SNP from the output of samtools mpileup at SNPs in the 1000 Genomes Project strict mask. BAF per bin was calculated by summing reads supporting alternate and reference alleles and taking into account phase information. Calling was conducted using 100 kb bins.

Following segmentation, raw CNA calls were filtered to identify high confidence events. The filtering thresholds for copy number and BAF were adjusted to capture integer copy number changes expected in a single cell. We applied the following criteria:

- Deletions: Segments were classified as deletions if they exhibited a normalized copy number below 1.5 and a strong BAF imbalance (|delta_BAF| > 0.4), indicating the complete loss of one allele.
- Duplications: Duplications were defined by an increased copy number (CN > 2.5) and a BAF deviation |delta_BAF| within the range of 0.08 to 0.24 (for copy number 3, the expected value is 1/6).
- CN-LOH: These events were identified in segments with a diploid copy number state (1.6 < CN < 2.4) that simultaneously showed a strong allelic imbalance (|delta_BAF| > 0.4).

To ensure the BAF signal was robust, all reported events were required to span at least 10 bins of heterozygous SNPs. Furthermore, we restricted our analysis to large-scale events by applying a minimum size filter of 10 Mb.

As a result of QC, four cells were deemed outliers and excluded from the analyses. We propose two possible explanations for their CNA profiles. The widespread chromosomal losses—some of which were homozygous (**Figure S6**)—may indicate nuclei with incomplete chromosome sets. This could result from chromosomes segregating into micronuclei and being lost during nuclear sorting. In cells exhibiting multiple smaller deletions, most of them span megabase-scale regions, far exceeding the typical size of PTA-amplified DNA fragments. Moreover, these deletions are not randomly distributed across the genome; rather, they are concentrated on specific subsets of chromosomes within each cell (**Figure S6**). We hypothesize that these regions may be selectively protected from amplification. The mechanism behind such a protection remains unclear. One possibility is that these deletions reflect a biological feature of senescent cells—specifically, that the protected regions correspond to senescence-associated heterochromatin foci (SAHF)^52^, which may inhibit access of polymerase during the PTA reaction. Cells showing losses of multiple chromosomes may represent an extreme manifestation of this phenomenon, with entire chromosomes being sequestered into SAHF and thereby excluded from amplification.

#### Analysis of aneuploidies on chromosome Y

Based on read depth, we identified 7 cells with Y loss (including one of the four outliers in **Figure S6**) and 2 with Y gain (**Figures S7A and S7B**). We used the seven cells with Y-loss for haplotype phasing of heterozygous SNPs in the pseudoautosomal region (PAR1). Because these cells retained only the X chromosome, the allele observed with zero count at heterozygous sites in PAR1 directly defines the Y-chromosome haplotype. We observed a high degree of concordance in the monoallelic pattern of these sites across all seven cells (**Figures S7C and S7D**), which allowed us to phase with high-confidence using majority voting. We then applied this phasing to high-quality heterozygous SNPs (within the 1000 Genomes Project strict mask) to analyze haplotype frequency distributions across all samples. This analysis confirmed our initial read depth-based classifications. In the seven cells with Y-loss, the haplotype frequencies for heterozygous sites collapsed to 0, indicating the complete absence of the Y-linked haplotype. Conversely, the two cells Y- gain exhibited a frequency peak shifted to approximately 2/3, a signature consistent with a trisomic XYY state. The other cells showed the expected diploid peak centered at 1/2 (**Figure S7B**).

#### Junction-spanning read analysis with Manta

Manta^43^ was run with default parameters, comparing each single-cell sample against the matched bulk sample to identify somatic structural variants (SVs). We first filtered Manta’s output for deletions and duplications larger than 100 kb. These initial calls were treated as candidates and subsequently validated using CNVpytor, requiring evidence of a corresponding integer-level copy number change and a significant BAF shift within the region. This stringent, two-step approach yielded 22 confident calls. Of these, 16 were located in T-cell receptor (TCR) loci, consistent with V(D)J recombination, while the remaining six are detailed in (**Figure S11**).

#### Reconstructing rearrangements in cell WashU_Lu_S2

For the large, sub-chromosomal CNAs previously identified by CNVpytor, we used Manta to provide orthogonal, breakpoint-level support. We first confirmed that these large events did not overlap with Manta’s deletion and duplication calls, indicating their complex nature. We then analyzed all break-end (BND) Manta’s calls ignoring Manta default filters. For cell WashU_Lu_S2, this approach successfully identified supporting BND calls for all predicted breakpoints (**Figure S9**), confirming the complex rearrangement model (**Figure 5D**).

Remarkably, just a few additional large or inter-chromosomal rearrangements were identified that were not supported by CNV analysis, indicating remarkable match between the orthogonal evidence (i.e., read depth, BAF, and discordant reads) and excluding the possibility of amplification induced error. Similar to analysis in PAR1 regions, we phased SNPs on chromosome 2 using an outlier cell (**Figure S6**) and uncovered that the deletion and the duplication in WashU_Lu_S2 cell occurred on different haplotypes. Fusion between chromosomes 7 and 17 was not called by Manta but was identified by investigation of split reads in IGV (**Figure S9**).

#### Applying HiScanner

HiScanner^41^ was run on all PTA BAMs and used the raw GATK output table (`gatk/hc_raw.mmq60.bcf`) and Eagle-phased germline SNPs (`eagle/phase_filtered.bcf`) created by SCAN2 for BAF and phasing information. Additional HiScanner parameters were `rdr_only=false`, enabling the use of BAF alongside read depth signals, `binsize=500kb`, `lambda=512`, multi-sample segmentation=false, and maximum whole-genome doublings (`max_wgd=1`) to exclude whole-genome doublings. The HiScanner output (`final_calls/*`) was filtered to include only autosomes. All diploid CN segments were examined for copy neutral LOH (CN-LOH) by segmenting into 10-bin (∼5Mb) intervals and intervals with mean BAF < 0.25 were determined to be CN-LOH. To smooth out small CNAs that may represent noise, segments with NBIN ≤ 20 (i.e., length ≤ ∼10Mb) had their CN set to the first NBIN > 20 segment to its left if such existed, and if not to the first NBIN > 20 segment on its right. If the smoothing process created adjacent segments with equal CN, the segments were merged. To analyze sex chromosomes, HiScanner was run again in read depth-only mode (`rdr_only=true`) with otherwise the same parameters, over all chromosomes. RDR-only CNVs in the sex chromosomes were combined with the autosomal CNV call set only when classifying cells as aneuploid.

## Notes

### Competing Interest Statement

Peter Park a member of the Scientific Advisory Board for Bioskryb Genomics and holder of software license for xTea (transposable element detection). Ji Won Oh is the founder and CEO of Absolute DNA, with no direct relation to this study, and the interests are managed by University-Industry Foundation in Yonsei University Health System in accordance with their conflict-of-interest policies; Dan Landau is on scientific Advisory Board of Mission Bio, Veracyte, Ultima, BioSkryb and has received prior research funding from 10x Genomics, Ultima Genomics, Oxford Nanopore Technologies and Illumina. He has multiple patents on duplex and single-cell sequencing; C.A.W. is a paid consultant (cash, no equity) to Third Rock Ventures and Flagship Pioneering (cash, no equity) and is on the Clinical Advisory Board (cash and equity) of Maze Therapeutics. No research support is received.

