## Supplementary Tables, Figures and Methods for "A comprehensive view of somatic mosaicism by single-cell DNA analysis"

**This file contains:**

Supplementary table S1, S3-S4.

Supplementary figures S1 to S11.

|  | **Yale-BCM** | **BCH-Broad** | **Cornell-NYGC** | **Mayo-WashU** | | | | **Yonsei** |
| --- | --- | --- | --- | --- | --- | --- | --- | --- |
| **Sample** | Core | Homogenate | Core | Core | | | | Core |
| **Sample dissociation** | Dounce homogenizer | Dounce homogenizer | Singulator 100 | Dounce homogenizer | | | | Dounce homogenizer |
| **Nuclei extraction** | Sucrose | Sucrose (modified) | Sucrose | Minute^TM^ detergent-free NP40 buffer | | | | Minute^TM^ detergent-free NP40 buffer |
| **DNA dye** | N/A | DAPI | PI | AOPI | N/A | SYT09 | N/A | AOPI |
| **Lysis** | On ice | On ice | On ice | RT | Ice | Ice | Ice | RT |
| **PTA kit version** | v1 | v1 | v1 | v1 | v2 | v2 | v1 | v1 |
| **QC (initial)** | 4-loci PCR | 4-loci PCR | - | 4-loci PCR | | | | 4-loci PCR |
| **Lib-prep** | PCR based | PCR free | PCR free | PCR based | | | | PCR free |
| **Lib-prep kit** | KAPA HyperPrep | KAPA HyperPrep | TruSeq | KAPA HyperPrep | | | | TruSeq |
| **Coverage** | 20-60X | ~30X | ~30X | ~30X | | | | ~30X |
| **Amplification site** | Yale | BCH | Weill Cornell | Yonsei | | | | Yonsei |
| **Sequencing site** | BCM | Broad | NYGC | WashU-VAI | | | | Theragen Bio |
| **# of cells** | 27 | 20 | 7 | 40 | | | | 8 |

**Table S1. Summary of experimental conditions across datasets.**

**Table S2 (external file). Data and analysis summary for all single cells.** Each cell is annotated with sample ID(s), dataset, tissue origin, sequencing depth, passed and failed QC metrics, numbers of SNV, indel and DNVs identified by SCAN2, inferred SNV and indel rates, inferred mutational signature exposures, whether the cell was identified as a T-cell based on rearrangement of the T-cell receptor loci, and whether the cell gained or lost a copy of chrY. BCH: Boston Children’s Hospital.

| Chr | Location | Forward | Reverse | Amplicon Size (bp) |
| --- | --- | --- | --- | --- |
| Chr 5 | 108698573- 108698712 | GGAGTCATCCTCCAGGTTATTGTTACCATC | CCTTGGAAGAGGGAGAAATTCCTTGGTTA | 140 |
| Chr10 | 117704859- 117705381 | CTTTCCGCCTAACTAGAATGCAGACCA | CGCTCGTGTTGGGAAGAAGACTCC | 523 |
| Chr15 | 42117570- 42117992 | TGCTGGAGCAATACTCAGAACTGTTGC | GCTAATCCCTGCAGTAATTTCAAATGGCT | 423 |
| Chr20 | 17970668- 17970958 | CTGGACCAAGTGGCTTCTTCGACTAG | GCGTGCCGAAGTCTAGGTCTTTATATCTAG | 291 |

**Table S3. Primer sequence for 4 loci PCR.**

| **Chromosome** | **Start** | **End** |
| --- | --- | --- |
| chr1 | 125162649 | 125189049 |
| chr1 | 143184000 | 143276000 |
| chr2 | 32915000 | 32917000 |
| chr2 | 90375650 | 90404200 |
| chr2 | 239865880 | 239869330 |
| chr3 | 93450000 | 93471000 |
| chr4 | 49095199 | 49161499 |
| chr4 | 190101449 | 190118199 |
| chr4 | 190172399 | 190186749 |
| chr5 | 3321225 | 3325000 |
| chr5 | 49594997 | 49608047 |
| chr5 | 49625797 | 49636197 |
| chr5 | 49641047 | 49672347 |
| chr5 | 139445999 | 139459549 |
| chr6 | 34002999 | 34013099 |
| chr6 | 34039099 | 34049999 |
| chr6 | 34065799 | 34078699 |
| chr7 | 100947349 | 100964499 |
| chr7 | 100986049 | 101004999 |
| chr7 | 158128500 | 158133500 |
| chr10 | 42061550 | 42105000 |
| chr13 | 30837149 | 30849999 |
| chr13 | 35756249 | 35768749 |
| chr16 | 34571000 | 34596500 |
| chr16 | 45000000 | 46405000 |
| chr18 | 0 | 113000 |
| chr20 | 31047050 | 31080900 |
| chrX | 10001 | 2781479 |
| chrX | 155701383 | 156040895 |
| chrY | 10001 | 2781479 |
| chrY | 11295049 | 11339249 |
| chrY | 56887903 | 57227415 |

**Table S4. Regions excluded from SNV, indel and DNV analysis by SCAN2.** Regions with anomalously high read depth and the pseudo-autosomal regions were not analyzed for small mutations with SCAN2.

**Supplementary Methods**

**Multiplex PCR conditions:** PCR reactions were performed in 20 µl volumes containing 1–10 ng of template DNA (either genomic or WGA) and 0.2–0.5 µM of each primer. Thermal cycling was carried out with an initial denaturation at 95 °C for 2 min, followed by 40 cycles of annealing (95 °C for 10 s) and extension (60 °C for 30 s). Fluorescence acquisition was performed during the annealing/extension steps. PCR products were analyzed by agarose gel electrophoresis to verify specific amplification and expected fragment sizes.

Successful amplification of all four loci confirmed the presence and integrity of representative genomic regions and served as a quality control step before downstream sequencing. All primer pairs were first validated in singleplex format to assess specificity and eliminate primer-dimer formation, and multiplex conditions were optimized accordingly. Each run included both positive (gDNA) and negative (no template) controls.

**PCR Reaction Conditions**

| Component | Volume/reaction | Final concentration |
| --- | --- | --- |
| 10× PCR Buffer | 1 µl | 1× |
| MgCl₂ | 0.6 µl | 1.5 mM |
| dNTP mix (10 mM of each) | 0.5 µl | 400 µM of each dNTP |
| Primer cocktail | 2 µl | 0.5 µM each |
| HotStarTaq DNA Polymerase | 0.1 µl | 0.5 units/reaction |
| Distilled water | Variable | – |
| Template DNA (added at step 4) | 20 ng | ≤1 µg/reaction |
| Total reaction volume | 10 µl |  |

Thermal cycling was performed using a touchdown PCR protocol consisting of an initial denaturation at 94 °C for 15 min; followed by 12 cycles of denaturation at 94 °C for 1 min, annealing at 68 °C for 1 min with a decrement of 1 °C per cycle, and extension at 72 °C for 1 min; then 26 amplification cycles at 92 °C for 1 min, 55 °C for 1 min, and 72 °C for 1 min; followed by a final extension at 72 °C for 10 min.

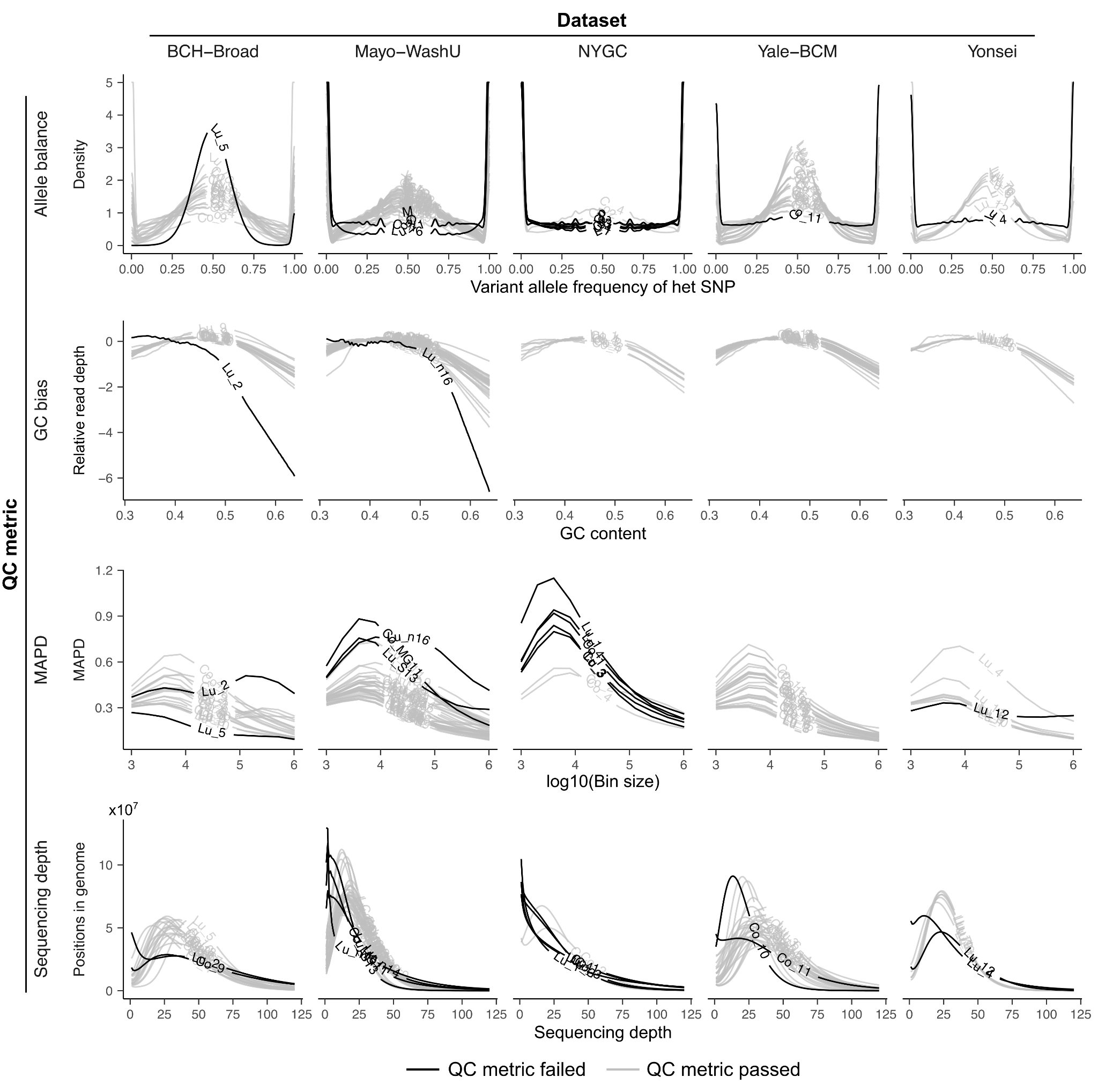

**Figure S1. Quality control metrics and visual identification of outlier cells.** QC failures were identified manually by comparison to all other cells across datasets. Each curve in each plot represents one single cell, which is identified by a shortened alias (**Table S2**); dark lines indicate QC failure of the specified cell for the particular QC metric plotted. Allele balance measures evenness of amplification between the maternal and paternal alleles by aggregating the VAF measured in each single cell at heterozygous SNP sites identified in bulk; concentration near ½ indicates better allele balance. GC bias reveals how amplification is affected by the GC content of the DNA being amplified; flatter curves indicate less bias. MAPD measures amplification variability by comparing the difference in sequencing depths of neighboring bins across the genome; lower MAPD indicates more even amplification. Poor sequencing depth is indicated by large fractions of the genome at very low depth, often lacking a mode near the target sequencing depth or by large fractions of the genome being sequenced at very high depth, leading to a “fat” right tail and depressed mode.

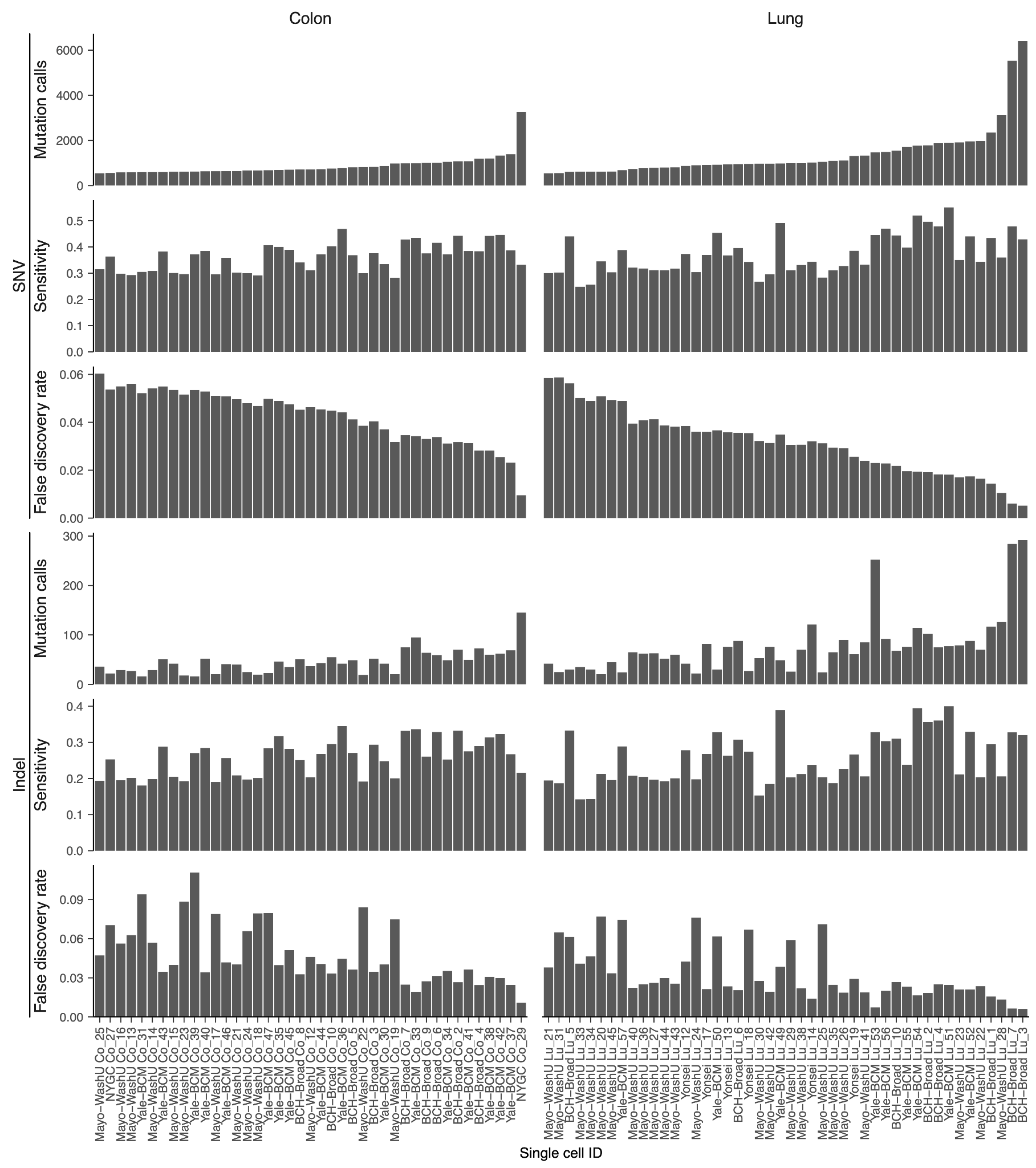

**Figure S2. Sensitivity and false discovery rates for SCAN2 somatic SNV and indel analysis.** Total number of mutation calls (top), sensitivity estimated by germline variant analysis (middle) and false discovery rate (bottom) for each single cell. Cells are ordered by total SNV calls.

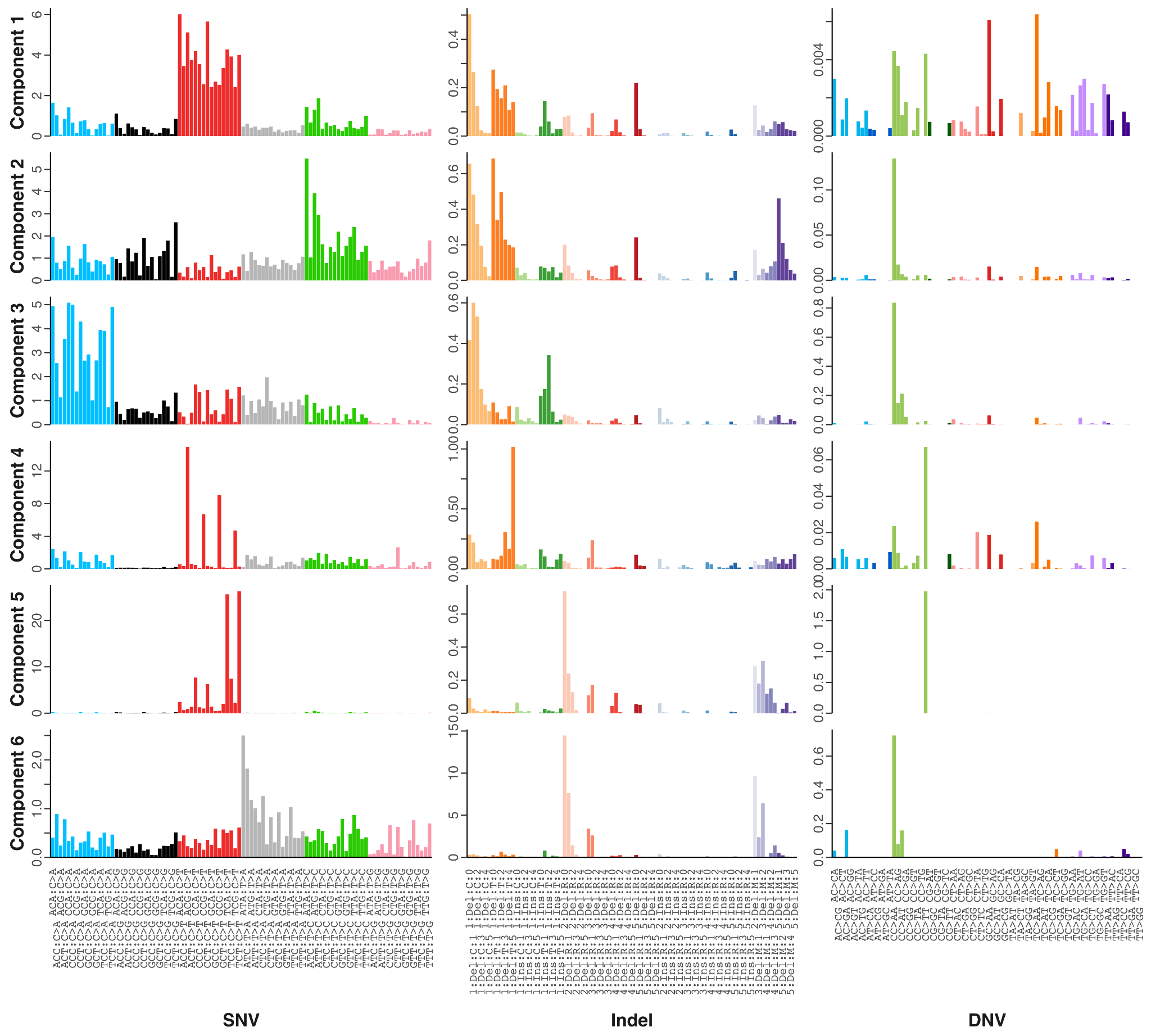

**Figure S3. *De novo* decomposition of mutation spectra into 257-dimensional components.** Unlike standard signature analysis in which mutation types such as SNVs and indels are analyzed separately, we combined SNVs, indels and DNVs into joint 257-dimensional vectors for each cell. This resulted in 6 components. y-axis values are the percentage represented by a channel among all 257 mutation channels.

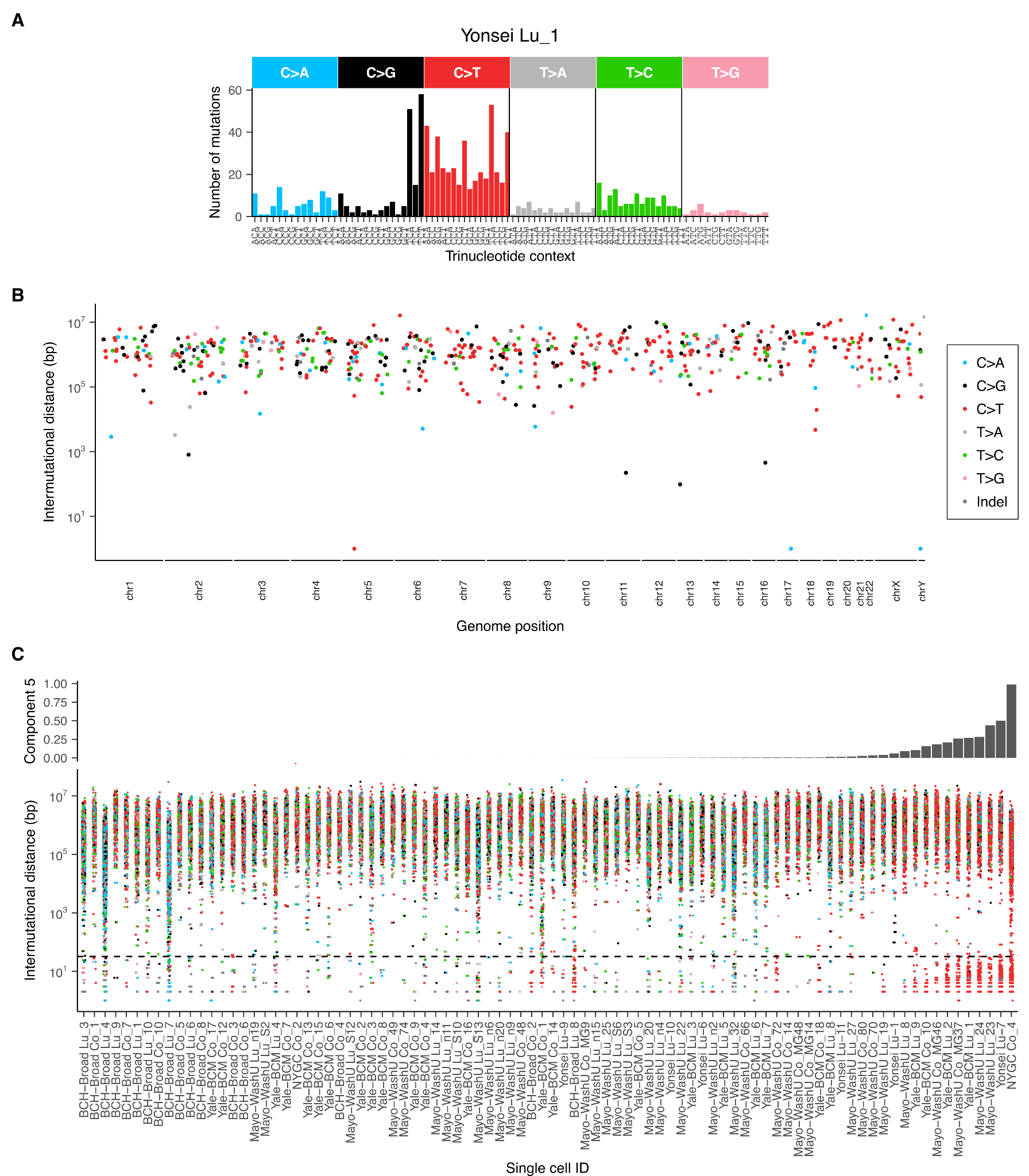

**Figure S4. Lung cell exhibiting both APOBEC COSMIC signatures SBS2 and SBS13 but no evidence of kataegis.**

**A.** 96-dimensional SBS spectrum for cell Yonsei Lu_1.

**B.** Rainfall plot showing distances between successive somatic SNVs and indels. DNVs are shown with intermutational distance=1. The absence of mutation clusters indicates kataegis, a common event associated with APOBEC mutagenesis, did not occur in this cell. Colors correspond to mutation types.

**C.** Intermutational distance distributions for all cells passing SNV and indel QC criteria. Each point represents the distance between two successive mutations. These plots omit genomic position, so cannot yield information about localized processes like kataegis, but are more compact and allow plotting of a large number of cells. Dashed line: intermutational distance = 32. Cells are ordered by component 5 (APOBEC-like) exposure. Colors are identical to panel **B**.

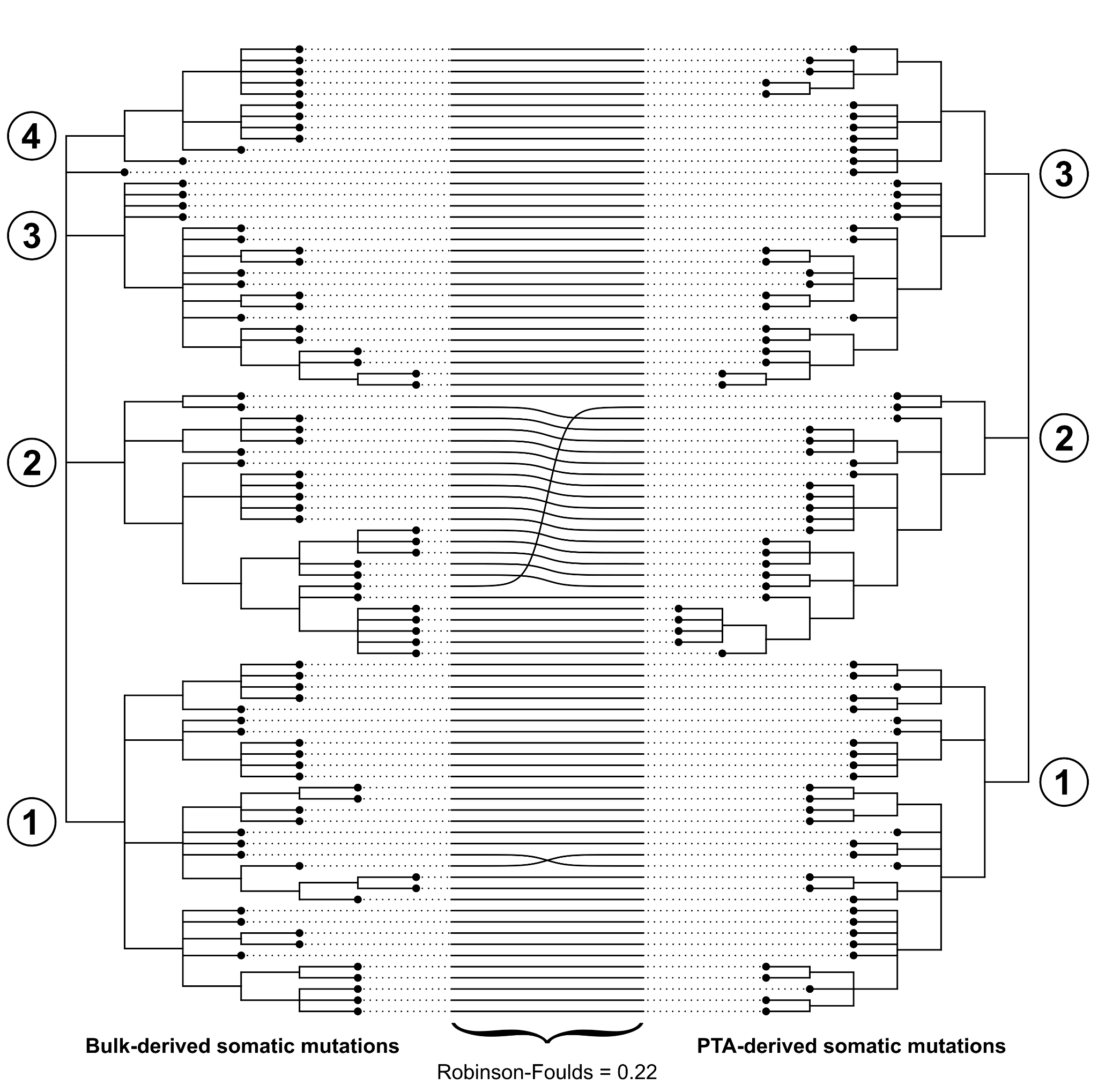

**Figure S5. Lineage trees reconstructed from mutations found in single cells and in bulk** **are nearly identical.** Lineage trees were reconstructed by applying Sequoia to two different sets of somatic mutations: those derived only from deep bulk sequencing and those derived only from PTA-amplified single cells. Major clades leading to the zygote are marked with numbered circles. The primary difference between the trees is the joining of clade 3 and 4 in bulk into a single clade 3 in the single-cell tree, which was caused by a mutation only called in single cells. Despite not being called, the somatic mutation was present in bulk.

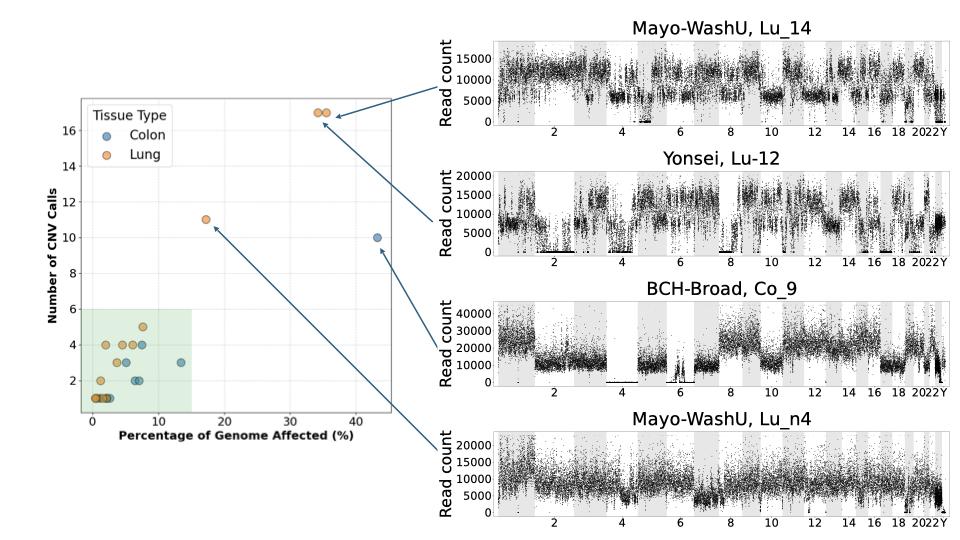

**Figure S6. Identification of outlier cells for CNV analysis.** Each cell, including the outliers identified in small mutation (SNV, indel, DNV) analysis, is plotted by its number of CNA calls and the proportion of the genome affected by a CNA. Four cells were deemed outliers and their read depth profiles are shown.

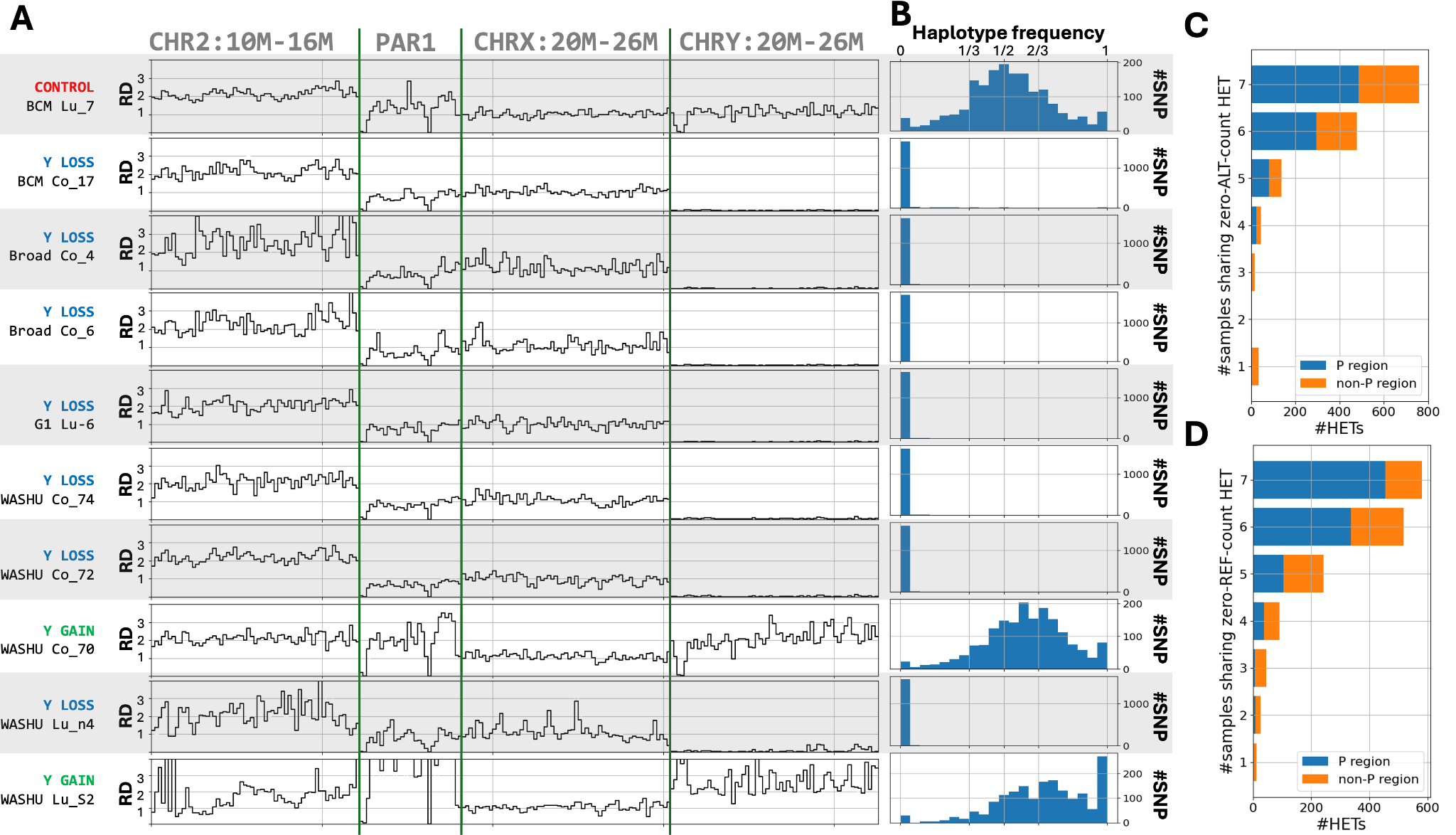

**Figure S7. Y chromosome aneuploidy in single cells. (A)** Read depth profiles across four genomic regions for a control cell, seven cells with Y chromosome loss (including one cell that failed CNA QC), and two cells with Y chromosome gain. The regions include a segment of an autosome, the pseudoautosomal region (PAR1), and non-PAR segments of chromosomes X and Y. Y-loss cells show a near-complete loss of read depth in the non-PAR Y region, while Y-gain cells show increased read depth in the Y and PAR1 regions. **(B)** Allele frequency distributions for heterozygous sites (HETs) located in the PAR1 region. In the control cell, allele frequencies cluster around 1/2. In Y-loss cells, allele frequencies are zero, while in Y-gain cells, allele frequencies are shifted to 2/3, which is characteristic of a trisomic state (e.g., XYY). **(C, D)** Analysis of consistent haplotype phasing across the seven Y-loss cells. The bar charts quantify the number of HETs that appear homozygous because only the alternate allele (C) or reference allele (D) is detected. The y-axis shows the number of cells that share this monoallelic pattern. The high number of HETs is consistently phased across all seven Y-loss cells.

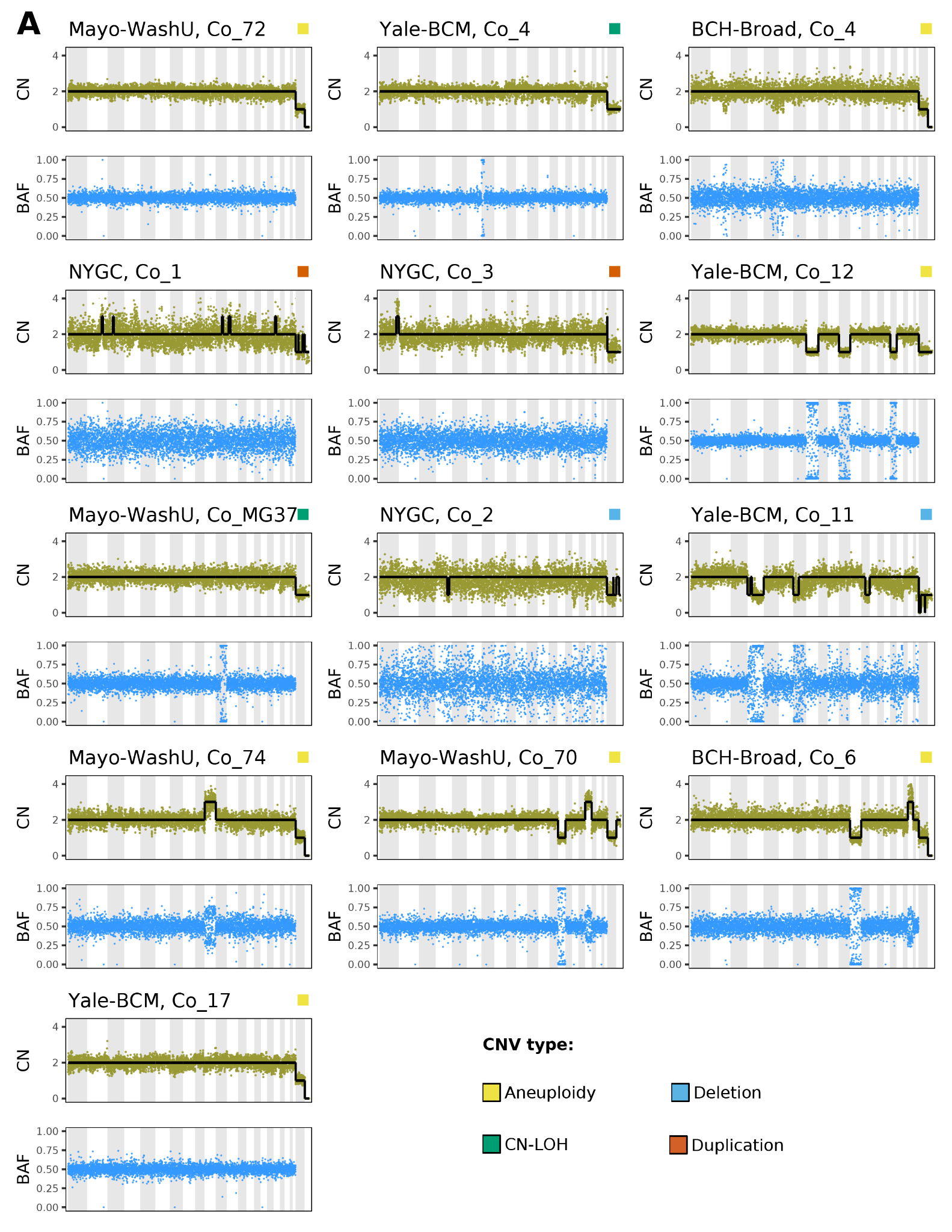

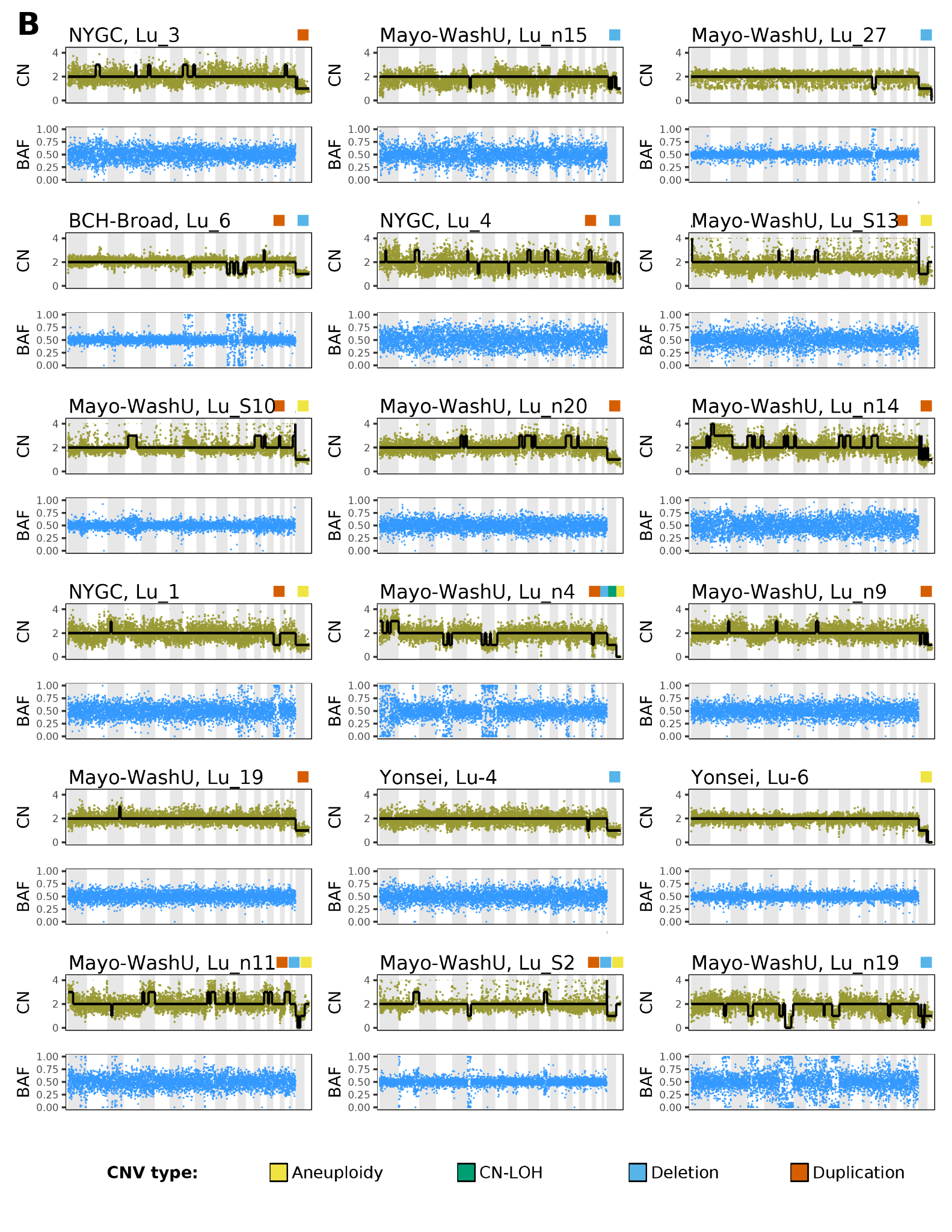

**Figure S8. All cells with chromosomal aneuploidies and large CNVs.**

**A.** Colon cells with at least one CNV.

**B.** Lung cells with at least one CNV.

Colored squares in the upper right corner of each cell plot denote which CNV types are present in that cell. Yellow: aneuploidy; Green: CN-LOH; Blue: sub-chromosomal deletion; Orange: sub-chromosomal duplication.

**A**

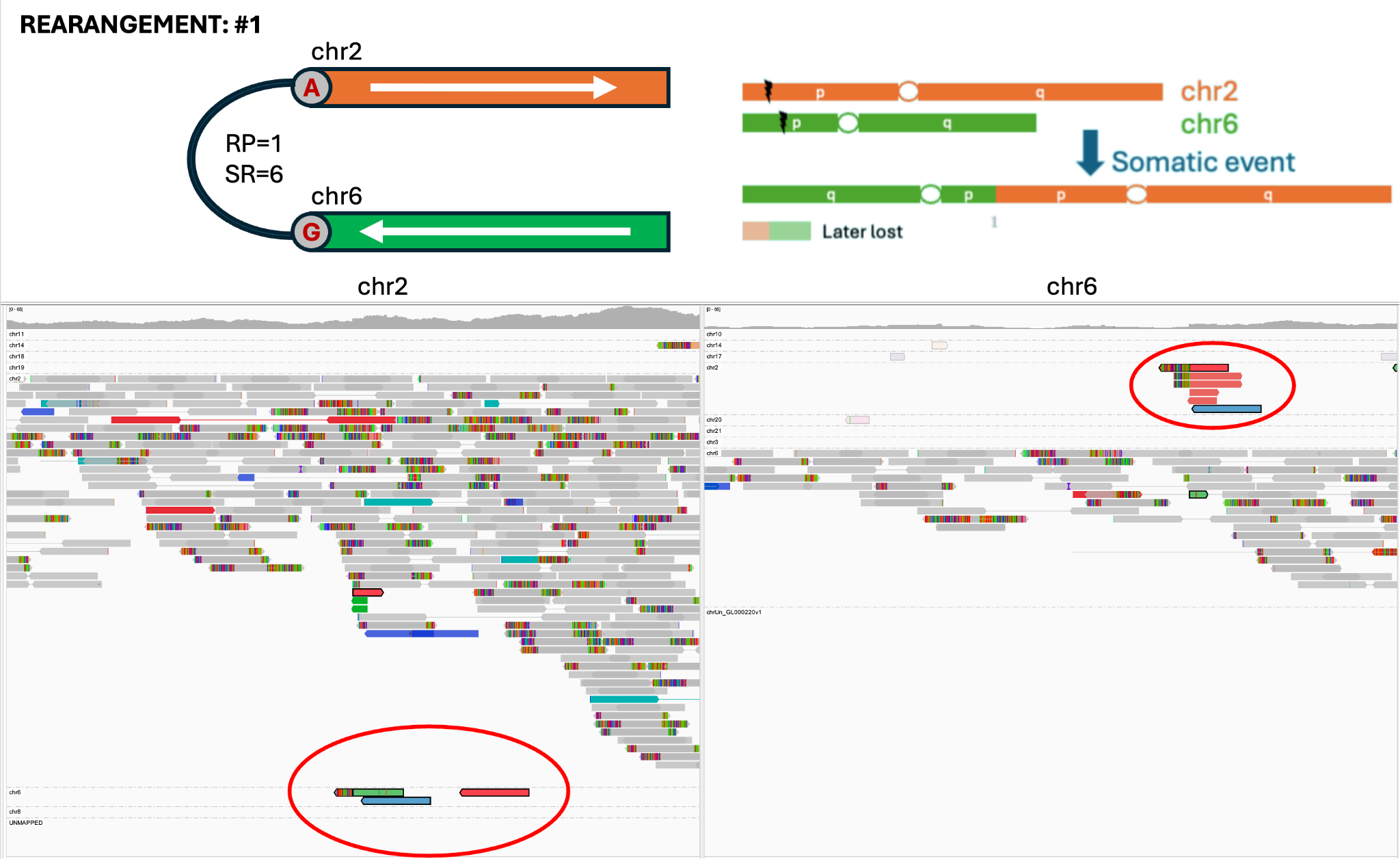

**B**

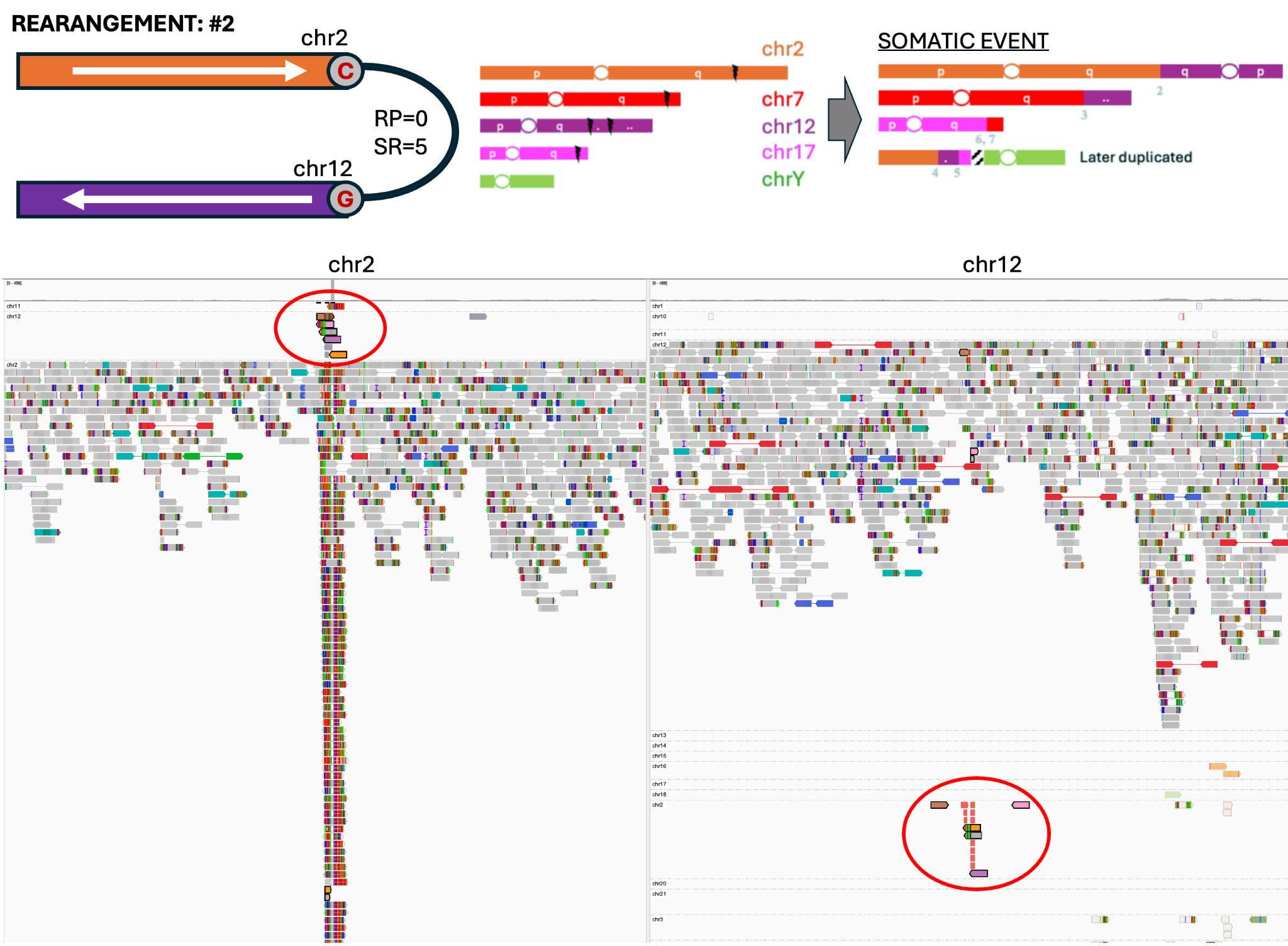

**C**

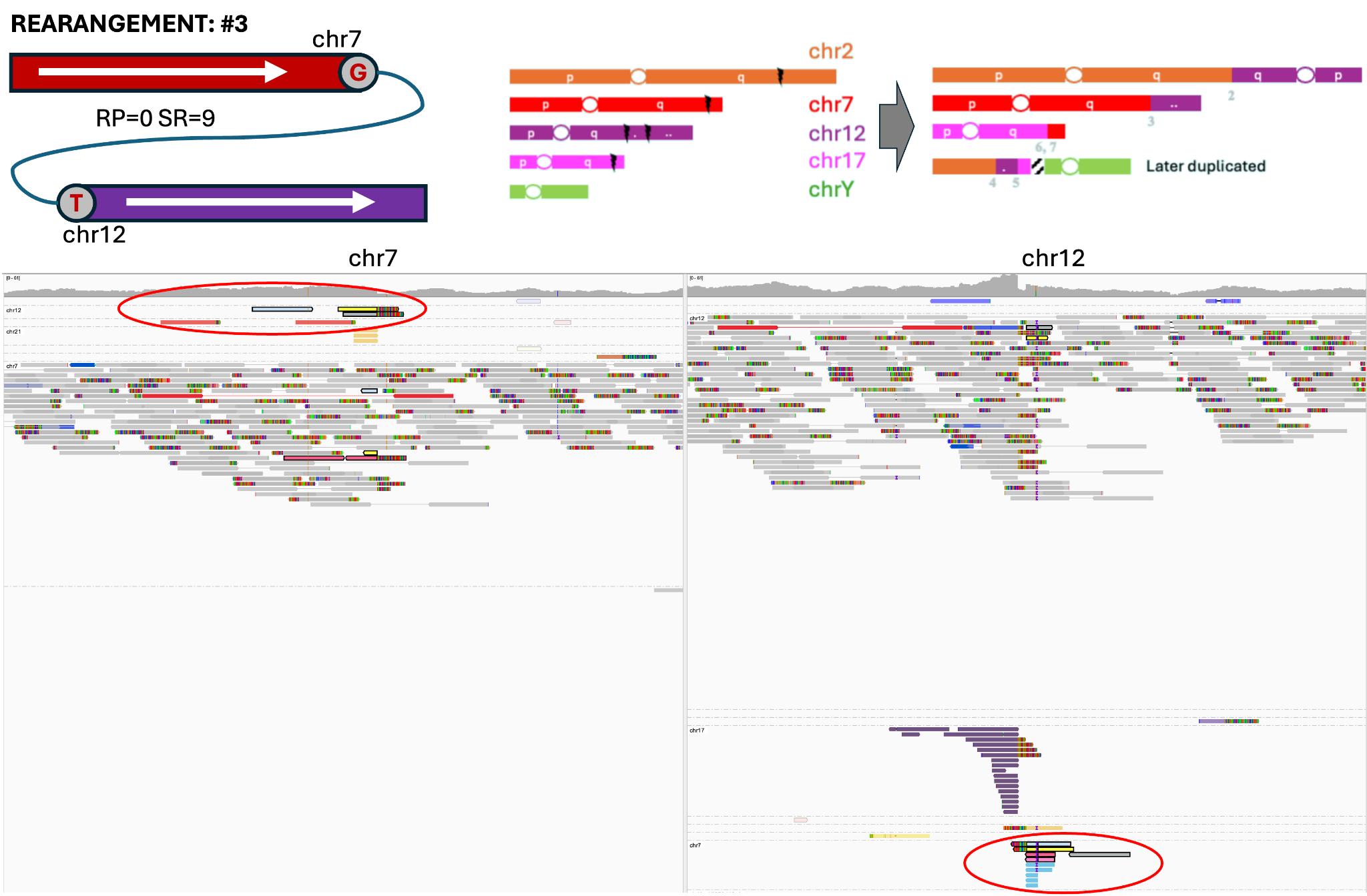

**D**

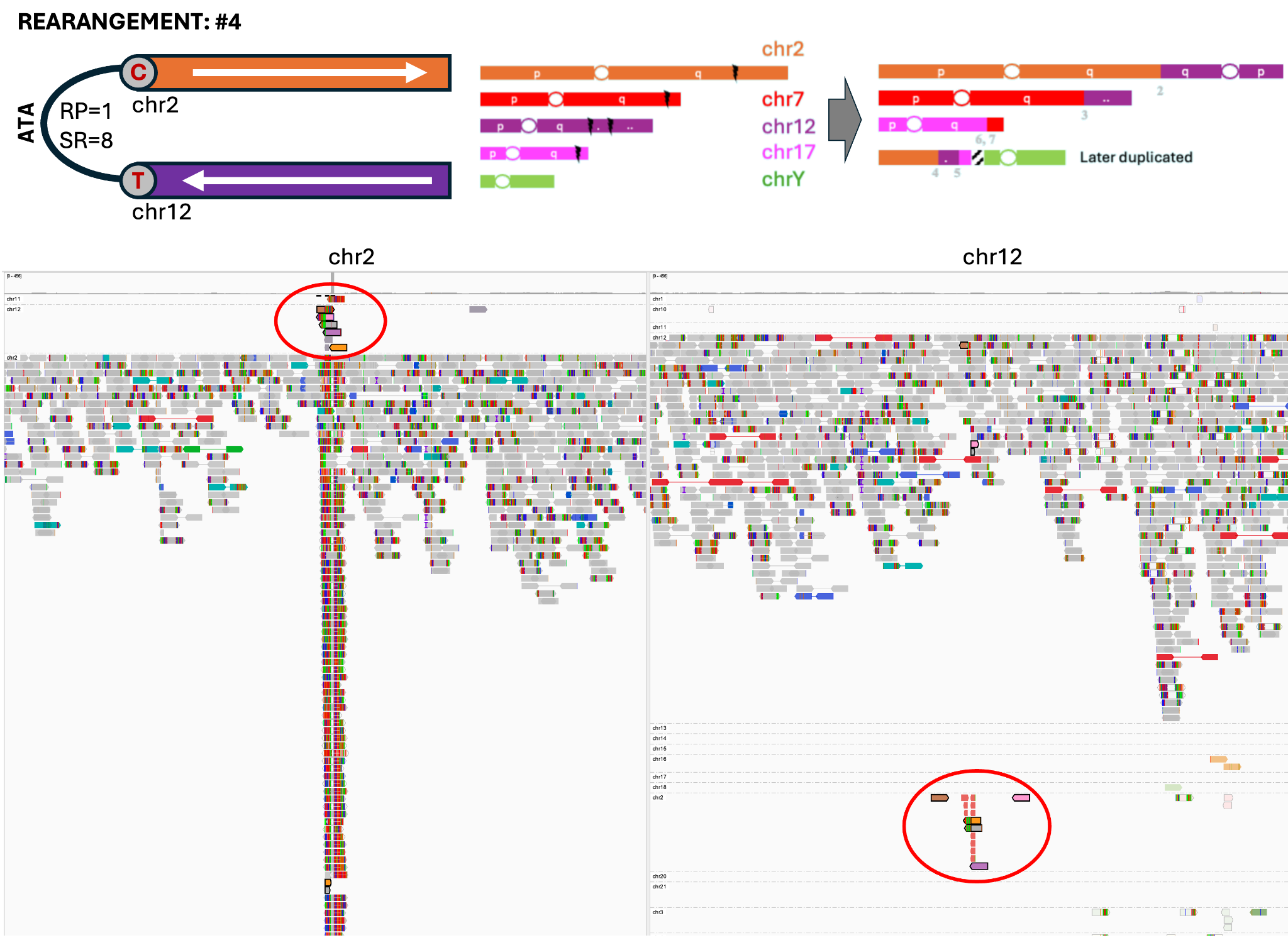

**E**

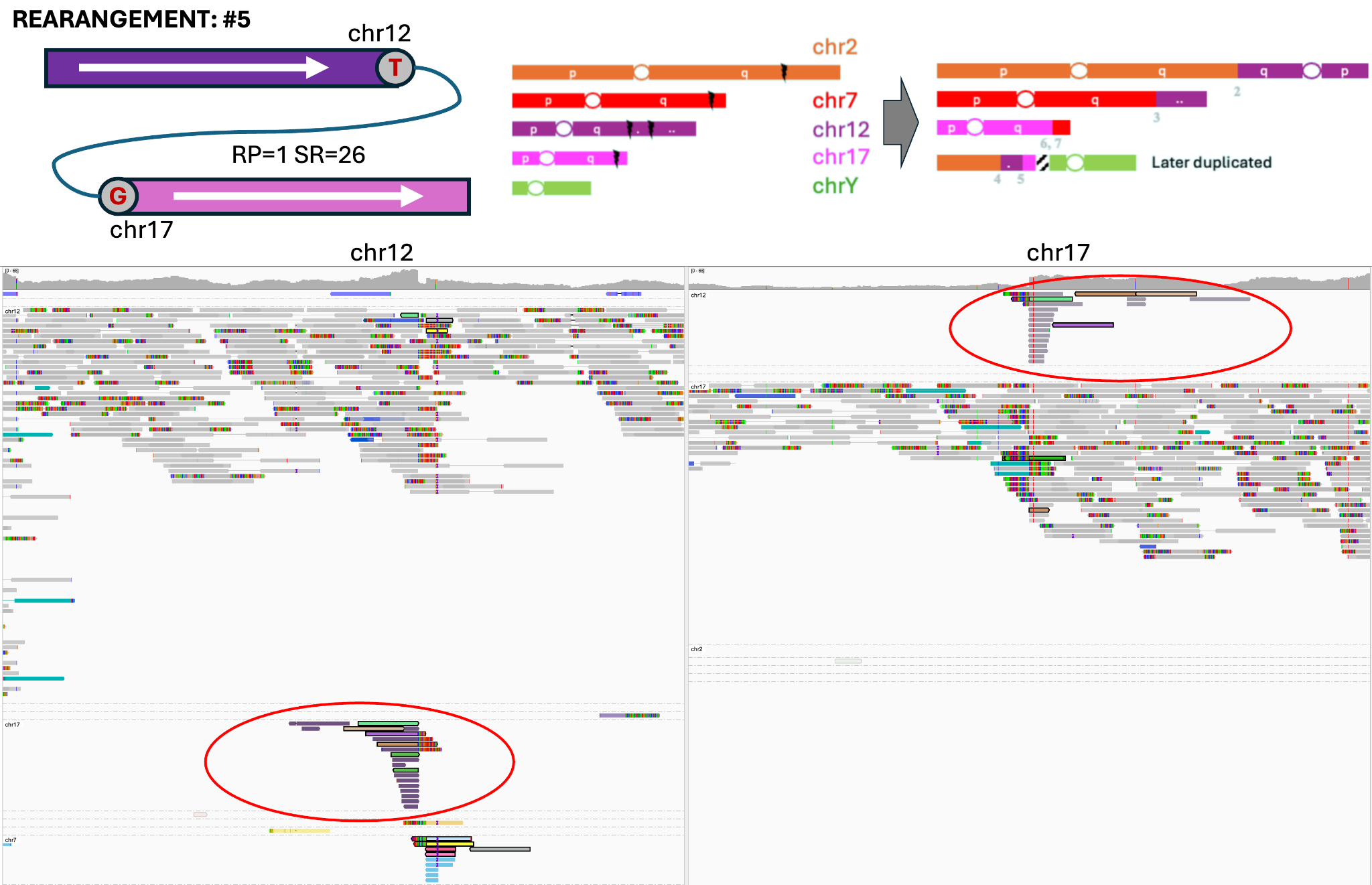

**F**

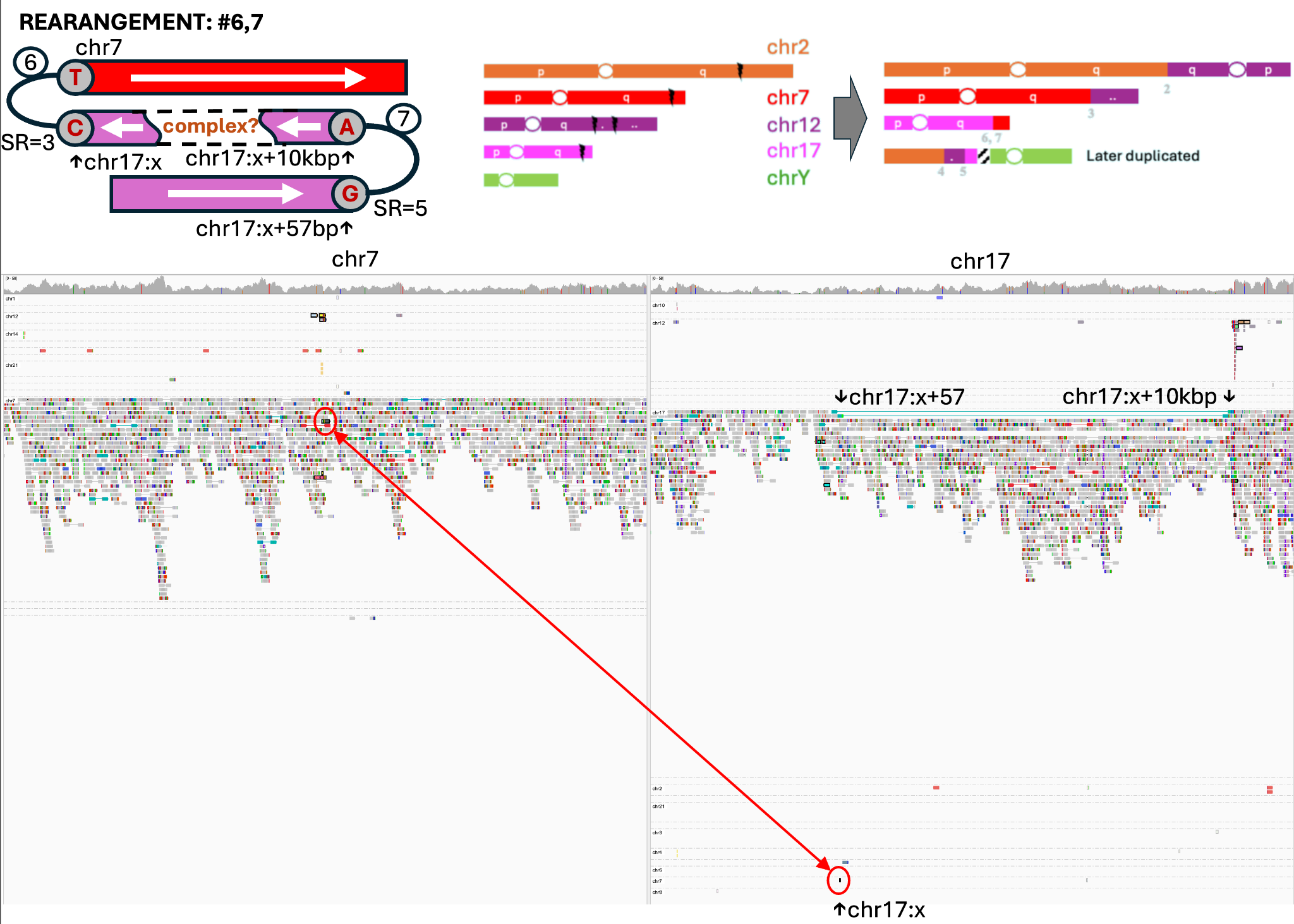

**Figure S9. Read support for the complex rearrangements observed in cell WashU, Lu_S2.** Panels **A–F** illustrate a rearrangement corresponding to the event shown in Figure 5D. Each panel combines a schematic diagram of rearrangement with an IGV snapshot. Genomic coordinates and explicit sequences have been removed in accordance with consent restrictions. In the IGV views, reads are grouped by chromosome of the mate and circled reads highlight supporting evidence for the rearrangement. The schematic diagrams indicate the chromosomal origin of joined fragments, their orientation (with reversed fragments shown on the left), number of discordant read pairs (RP) and split reads (SR) found by Manta, and the presence of microinsertion in the case of rearrangement #4.

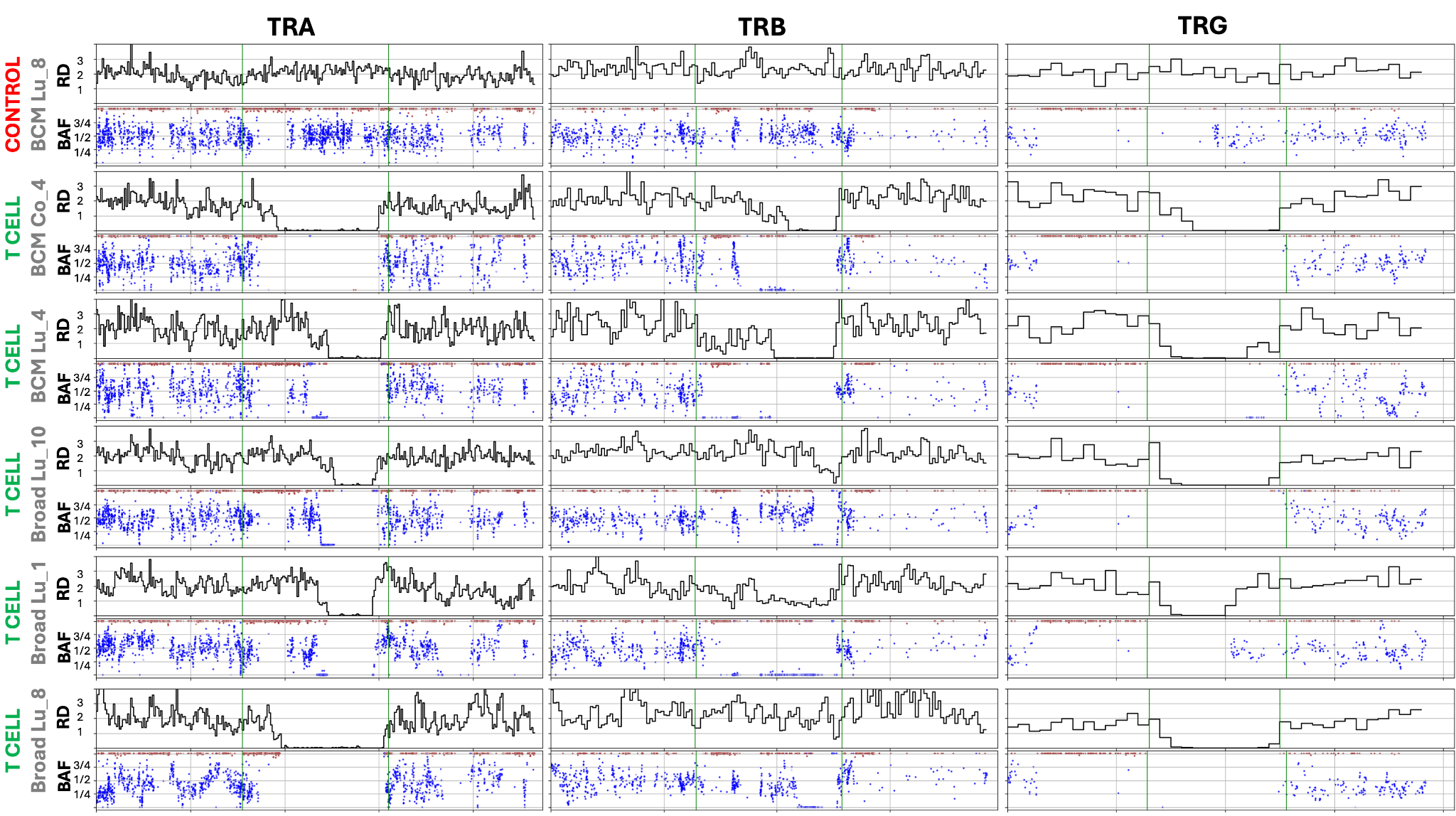

**Figure S10. Detailed plots of read depth and BAF at the T-cell receptor loci for the five identified T-cells.** RD: read depth normalized to ploidy; BAF: B-allele frequency of germline SNPs; TRA, TRB, TRG: T-receptor locus alpha, beta and gamma, respectively.

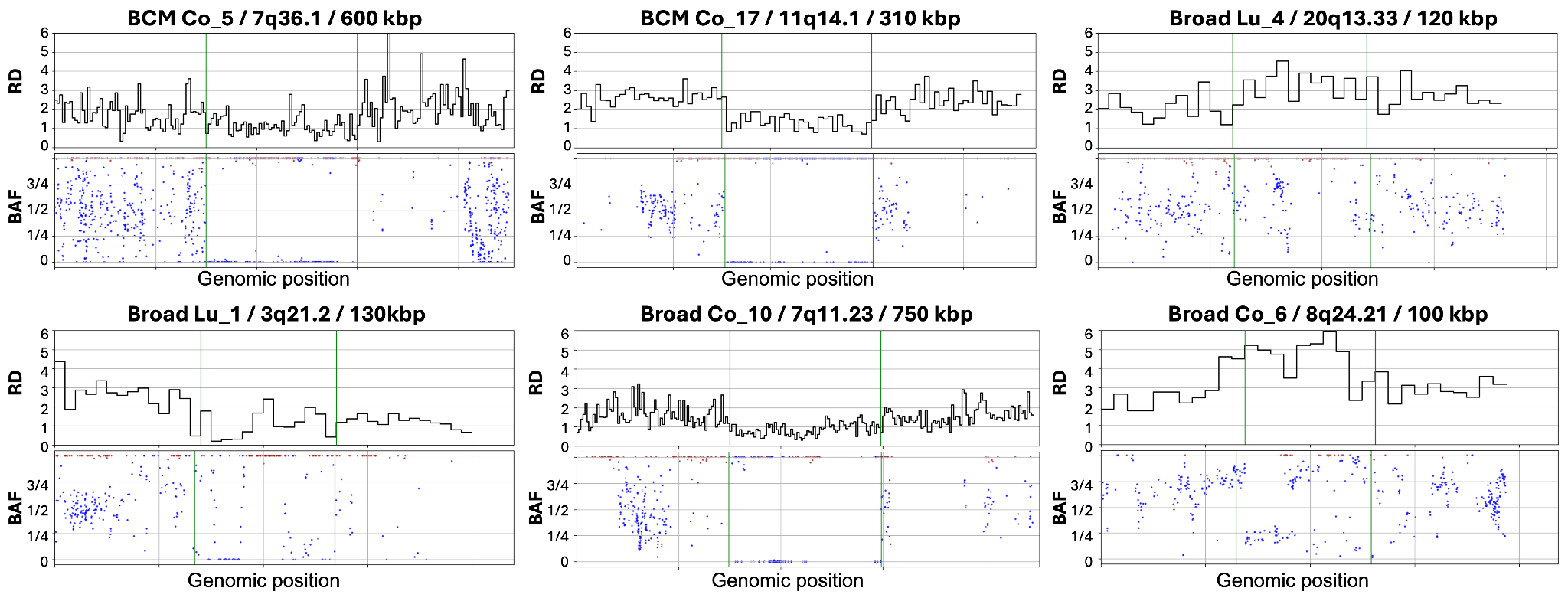

**Figure S11. Deletions and duplications with breakpoint-spanning reads outside of the T-cell receptor loci.** Panels include the sample name, the affected chromosomal region, and the size of the event rounded to 10 kbp. RD: read depth normalized to ploidy; BAF: B-allele frequency of germline SNPs;
